# Lupus-associated innate receptors drive extrafollicular evolution of autoreactive B cells

**DOI:** 10.1101/2024.01.09.574739

**Authors:** Danni Yi-Dan Zhu, Daniel P. Maurer, Carlos Castrillon, Yixiang Deng, Faez Amokrane Nait Mohamed, Minghe Ma, Aaron G. Schmidt, Daniel Lingwood, Michael C. Carroll

**Affiliations:** Program in Cellular and Molecular Medicine, Boston Children’s Hospital, Harvard Medical School, Boston, MA 02115, USA; Harvard Graduate Program in Virology, Boston, MA 02115, USA; Department of Microbiology, Harvard Medical School, Boston, MA 02115, USA; Department of Pediatrics, Harvard Medical School, Boston, MA 02115, USA; Ragon Institute of Mass General, MIT, and Harvard, Cambridge, MA 02139, USA; Department of Medicine, Emory University School of Medicine, Atlanta, GA 30307, USA

## Abstract

In systemic lupus erythematosus, recent findings highlight the extrafollicular (EF) pathway as prominent origin of autoantibody-secreting cells (ASCs). CD21^lo^CD11c^+^ B cells, associated with aging, infection, and autoimmunity, are contributors to autoreactive EF ASCs but have an obscure developmental trajectory. To study EF kinetics of autoreactive B cell in tissue, we adoptively transferred WT and gene knockout B cell populations into the 564Igi mice - an autoreactive host enriched with autoantigens and T cell help. Time-stamped analyses revealed TLR7 dependence in early escape of peripheral B cell tolerance and establishment of a pre-ASC division program. We propose CD21^lo^ cells as precursors to EF ASCs due to their elevated TLR7 sensitivity and proliferative nature. Blocking receptor function reversed CD21 loss and reduced effector cell generation, portraying CD21 as a differentiation initiator and a possible target for autoreactive B cell suppression. Repertoire analysis further delineated proto-autoreactive B cell selection and receptor evolution toward self-reactivity. This work elucidates receptor and clonal dynamics in EF development of autoreactive B cells, and establishes modular, native systems to probe mechanisms of autoreactivity.

## Introduction

Aberrant production of autoantibodies targeting nuclear self-antigens comprises an important disease hallmark of SLE^1-3^. The circulation and deposition of autoantibodies due to breach of B cell tolerance marks SLE serological activity and causes systemic organ inflammation^1,2^. Two primary pathways contribute to autoreactive antibody-secreting B cell (ASC) production: the germinal center (GC) and the extrafollicular (EF) pathways. The GC pathway leads to the persistent production of high- affinity, class-switched antibodies, whereas the EF pathway allows for a more rapid response, with B cells differentiating into ASCs outside the follicles^4-6^. In fact, SLE-associated human DN2 cells (CD11c^+^CD21^lo^T-bet^+^) and their so-called murine equivalent age-associated B cells (ABCs), are proposed to derive at least partly from the EF pathway and stand poised to differentiate into autoreactive ASCs ^4,7-1^^4^. Thus, the EF B cell differentiation pathway exhibits strong clinical relevance and generates pathogenic B cells as possible therapeutic targets.

While extensive phenotypical characterization of DN2-like B cell subsets has been performed in SLE patient-derived cells, our understanding of the mechanics behind these cells remains rudimentary. Human studies rely heavily on profiling the peripheral circulating population, and thus come with the inherent limitation of tissue representation. Furthermore, while human DN2 cells have been described as extrafollicularly-derived, much of this identity is derived from surface marker expression or from shared identities with infection-induced B cell subsets by previous studies^6,15^. Circulating immune cell profiling also does not support mechanistic studies, and murine-based tissue analyses remains crucial to define origin and temporal development of the pathogenic B cells. On the other hand, while ABC/DN2-like phenotypes were recapitulated in a number of lupus models, precise segregation between GC-dependent and -independent B cell development has been challenging^11,14^. In addition, a number of classic spontaneous lupus models were developed with a complex genetic background, and therefore bear intrinsic B cell anomalies that compromise the reliability of developmental studies^16,17^. Similarly, while the 564Igi-based mixed bone marrow chimera model was a powerful system for autoreactive GC profiling, it does not allow distinction between GC- and extrafollicular events^4^. Hence, a clean murine system with improved representation of the EF B cell differentiation pathway remains an imminent need to the field for comprehensive characterization of disease-associated subsets. While Ig transgenic mice might provide such system, their constrained BCR repertoire diversity and narrowed antibody specificity mask autoreactive B cell clonal evolution^12,18-2^^1^.

Innate receptors, including toll-like receptors and complement receptors, are vital for normal B cell development and demonstrate exacerbated importance in autoimmunity ^12,22,23^. Toll-like receptor 7 (TLR7) recognizes single-stranded RNA (ssRNA) and, upon engaging self-RNA, triggers intracellular signaling in activated B cells, providing a second signal that promotes escape of tolerance^24,25^. TLR7 expression is disease-associated^26,27^ and is required for murine ABC development and autoantibody production^11,14^. Additionally, it promotes the ex vivo differentiation of human DN2 cells and autoreactive ASCs with B cell receptor (BCR) co-engagement^8^. However, while TLR7 engagement is superficially understood to drive escape of tolerance by proto-autoreactive B cells, the receptor’s nuanced functions in the GC and EF differentiation pathways remain unaddressed *in vivo*^30^.

Interestingly, while DN2 cells/ABCs exhibit increased sensitivity to TLR7, they also downregulate the complement receptor CR2/CD21. With CD19, CD21 constitutes the BCR co-receptor complex that is crucial for lowering the threshold for B cell activation, thereby enhancing proliferation and differentiation^31-3^^3^. This receptor recognizes complement C3 fragments (iC3b, C3dg and C3d) and is capable of binding complement opsonized antigens^34,35^. Despite its role, the distinct association of CD21-low (CD21^lo^) cells with SLE and other chronic inflammatory diseases poses an apparent paradox. Therefore, refining the role of CD21 in early ASC differentiation is essential, and *in vivo* evidence remains limited largely due to the absence of an adequate mouse model.

We established an adoptive transfer mouse model system in which B cells with natural BCR repertoires were introduced into the 564Igi transgenic strain as the host^36^. 564Igi mice express a single immunoglobulin heavy and light chain insertion specific for nuclear antigen, creating a predominantly RNA-driven autoimmune environment with T cell help in which naïve B cells from non-autoimmune donor mice proliferate and differentiate into self-targeting ASCs^12^. The 564Igi host provides support for escape of tolerance by donor B cells, but does not directly interfere with the donor repertoire, thus preserving donor BCR diversity and clonal integrity (**Fig. 1a**). The donor cells can be analyzed from the time of transfer to days after, thus adding a temporal dimension that enables studying the rapid EF response.

**Figure 1.**
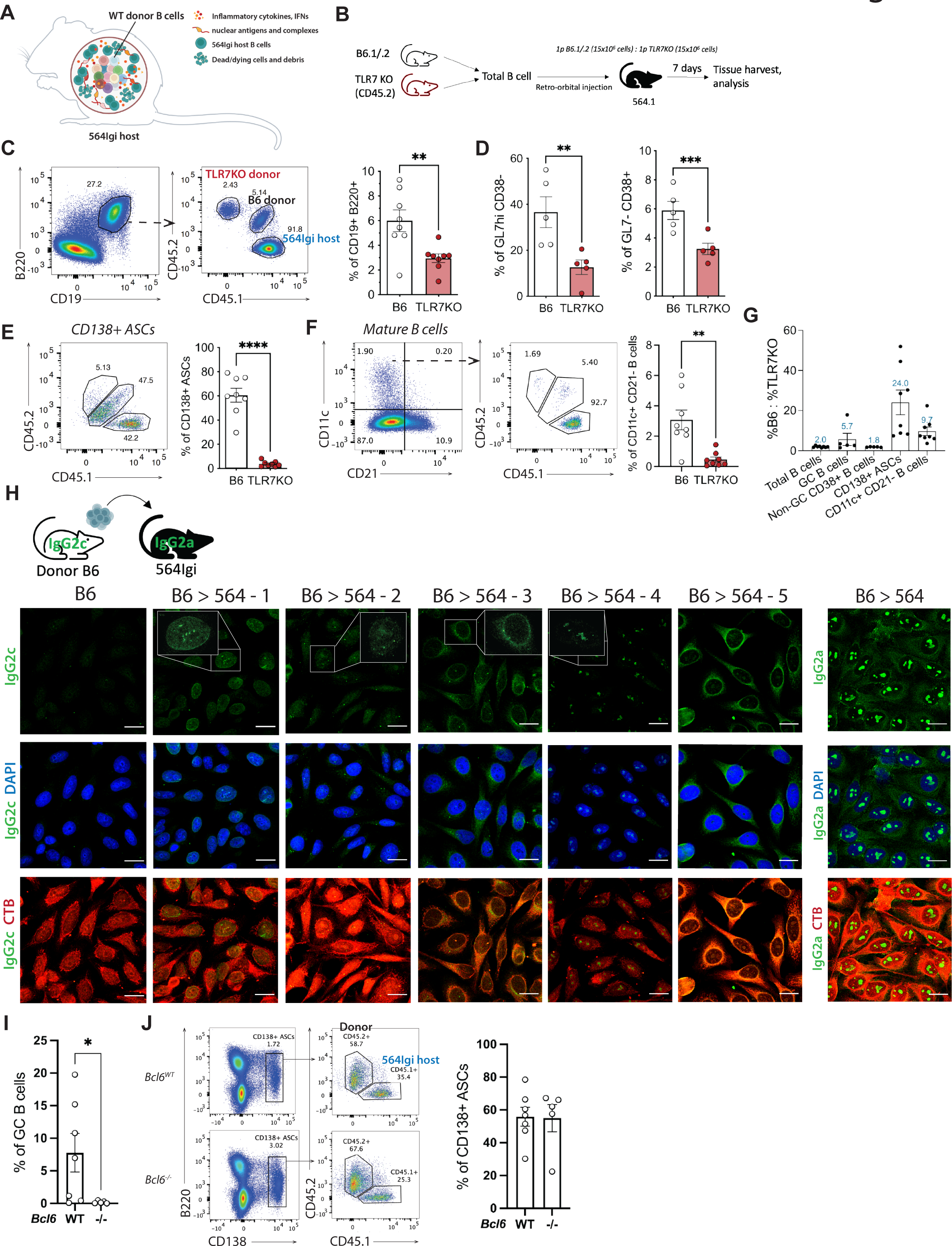
TLR7 drives efficient development of autoreactive B cells and EF ASCs with a DN2-like phenotype. (A) Schematic illustration of the adoptive transfer model. (B) Experimental set up of competitive WT/TLR7KO B cell adoptive transfer. (C) Representative flow cytometry plots and quantification of donor cell frequencies among total mature B cell compartment (CD19+ B220+) within competitive transfer 564Igi host (n=8). (D) Quantification of donor cell frequencies among GC (GL7hi CD38-) and non-GC mature (GL7- CD38+) B cells; gated from CD19+ B220+ B cells (n=5). (E, F) Representative flow cytometry plots and quantification of donor cell frequencies among CD138+ ASCs and CD11c+ CD21- CD138- B cells (n=8). (G) Ratio of %B6 to %TLR7KO cells across different B cell compartments from 7 days of transfer. (H) Representative images of HEp-2 cell immunofluorescence (IF) staining with adoptive transfer sera from 5 recipients and naive B6; CTB = cholera toxin B, cytoplasm; DAPI = nucleus. *B6>564-1*: nuclear dense fine speckled; *B6>564-2*: cytoplasmic/Golgi-like, *B6>564-3*: cytoplasmic discrete dots/GW body-like, *B6>564-4*: punctate nucleolar; *B6>564-5*: fine-speckled cytoplasmic. Images taken at 100x magnification, scale bars = 20 µm. (I) Quantification of Mb1cre^+/-^, (*Bcl6*^WT^, n=5) and Mb1cre^-/-^, Bcl6^fl/fl^ (*Bcl6*^-/-^, n=7) donor GC B cells (CD45.2+ GL7hi CD38-) among total GC B cells in 564Igi host at 7 days post-transfer. (J) Representative flow cytometry plots and quantification of Mb1cre^+/-^ , and Mb1cre^-/-^, Bcl6^fl/fl^ donor CD138+ ASCs. Statistical analysis with paired t test (competitive transfers) or Student’s t test; *=p<0.05, **=p<0.01; ***=p<0.001; ****=p<0.0001.

We utilize the adoptive transfer model to functionally delineate dynamics of TLR7 and CD21 on pathogenic B cells in their native tissue environment. We demonstrate that the two receptors function as molecular switches to modulate different aspects of donor cell tolerance breaking, division, and differentiation in the 564Igi host. Strikingly, we find that CD21^lo^ is a reversible phenotype while the receptor directly links to EF ASC differentiation. Tracking repertoire changes of naïve follicular B cells and their progeny subsets over time further elucidated evolution of proto-autoreactive B cells and demonstrated rapid selection and expansion of ASC clones with autoreactive features. Together, our findings advance current understanding of the fundamental biological circuitry behind EF autoantibody production and underscore the synergistic functions of innate receptors in driving the break of B cell tolerance.

## Results

### TLR7 drives the efficient development of autoreactive B cells and EF ASCs with a DN2-like phenotype

To characterize the role of TLR7 in driving escape of tolerance and ASC development, we adoptively transferred splenic B cells from B6 mice (CD45.1^+^CD45.2^+^, B6.1/.2) and *TLR7*-KO mice (CD45.2^+^) at a 1:1 ratio as competitors into a homozygous 564Igi host on a CD45.1 background (564.1). The congenic markers served as identifiers of the donor and host cells from different sources. Before transfer, we found that the naïve *TLR7*-KO splenic B cell compartment resembles that of the WT, except for a larger marginal zone B cell (MZB) compartment in aged mice (**Fig. s1A-D**). Spleens were harvested at 7 days post-transfer (d.p.t.), and *TLR7*-KO showed reduced generation of total mature (CD19^+^B220^+^), GC (GL7^hi^CD38^-^), and non-GC (GL7^-^CD38^+^) B cells (**Fig. 1B-D**). However, the largest reduction was in ASCs and DN2-like cells (CD11c^+^CD21^-^), which were respectively 24-fold and 9.7-fold reduced compared to those derived from WT cells (**Fig. 1E-G**). These results imply that, while TLR7 is not required for survival and maintenance, it is needed for efficient proliferation and differentiation of autoreactive donor B cells. Intriguingly, WT plasma cells (PCs) (CD138^+^B220^-^), but not plasmablasts (PBs) (CD138^+^B220^+^), also outcompeted the 564Igi host B cells by almost 40% although donor B cells were constrained in quantity (**Fig. s1E**, **F**).

To assess circulating autoantibodies, we exposed sera recovered from recipients at 7 d.p.t. to HEp-2 cells and observed varied IgG2c intracellular staining patterns. As the B6 heavy chain locus is derived from IgH-1b allotype, encoding IgG2c, whereas the 564Igi heavy chain encodes IgG2a, the antibody response indicates donor specificity against a diverse set of intracellular autoantigens^12,37^ (**Fig. 1H**). As controls, HEp-2 reactivity was absent from naïve B6 sera, and 564Igi anti-IgG2a staining showed strong nucleolar specificity as expected. Given the rapid emergence of ASCs and autoantibodies, we hypothesized they were produced through the EF pathway. To verify this, mb1-cre^+/-^,*Bcl6^fl/fl^* mice were generated with a B cell-specific deletion of *Bcl6*, which prevents B cells from differentiating into GC B cells^38^ (**Fig. 1I**). We transferred mb1-cre^+/-^,*Bcl6^fl/fl^* and mb1-cre^-/-^,*Bcl6^fl/fl^* B cells into separate 564Igi hosts and found no significant difference in ASC quantity, indicating that ASCs at 7 d.p.t. had likely matured via the EF pathway (**Fig. 1J**).

To examine if the accumulation of donor-derived DN2-like cells and ASCs was specific to autoimmune conditions, equal numbers of B6 (B6.1/.2) and *TLR7*-KO (CD45.2) B cells were adoptively transferred into naïve B6 CD45.1 (B6.1) mice (**Fig. s1I**). While both WT and *TLR7*-KO donor B cells were still present at 7 d.p.t., they made up a significantly lower proportion of the B cell compartment than in the 564Igi host (**Fig. s1J**). In B6 mice, the *TLR7*-KO B cells were approximately 50% less competitive than WT B cells and produced no ASCs (**Fig. s1J, K**). Lastly, few donor DN2-like cells were detected in the naïve host (**Fig. s1L**). Together, these results show that TLR7 is critical in supporting the rapid, extrafollicular development of autoreactive B cells.

### WT and *TLR7*-KO donor B cells occupy distinct splenic follicular niches and have divergent outcomes

To gain a better understanding of the developmental kinetics of donor B cells within the spleen, we adoptively transferred competing populations of B6.1/.2 and *TLR7*-KO B cells into a 564.1 host and harvested the spleens at 1, 4, 5, and 6 d.p.t. (**Fig. 2A**). We found that the WT B cells began to expanded at or before 4 d.p.t. (**Fig. 2B**) and again found that *TLR7*-KO had drastically reduced the number of DN2- like cells (**Fig. 2C**). Confocal immunofluorescent (IF) imaging revealed that, within the splenic white pulp, donor B cells entered and sparsely distributed across the B cell follicles at 1 d.p.t. At 4 d.p.t., the donor B cells packed close to the marginal zone (MZ) borders, as demarcated by CD169^+^ MZ macrophages. The donor cells began to spatially segregate at 5 d.p.t., with WT B cells occupying the follicular area and *TLR7*-KO B cells migrating to the outer MZ (**Fig. 2D, E**). Analysis of 32 white pulps from 5 d.p.t. spleens showed that the Mander’s Overlap Coefficient tM1, which represents the proportion of CD45.2 signals that overlaps with CD45.1, was significantly higher within follicles than within MZ, indicating preferential localization of WT B6.1/.2 B cells to the follicles and CD45.2^+^ *TLR7*-KO B cells to the MZ. This was likely the result of a contracted 564Igi MZ B cell compartment, leaving unfilled MZ space for the *TLR7*-KO cells to occupy^39^ (**Fig. s2A, B**). Together, the distinct splenic localization of donor WT and *TLR7*-KO B cells supports their diverging fates and may contribute to their contrasting developmental outcomes.

**Figure 2.**
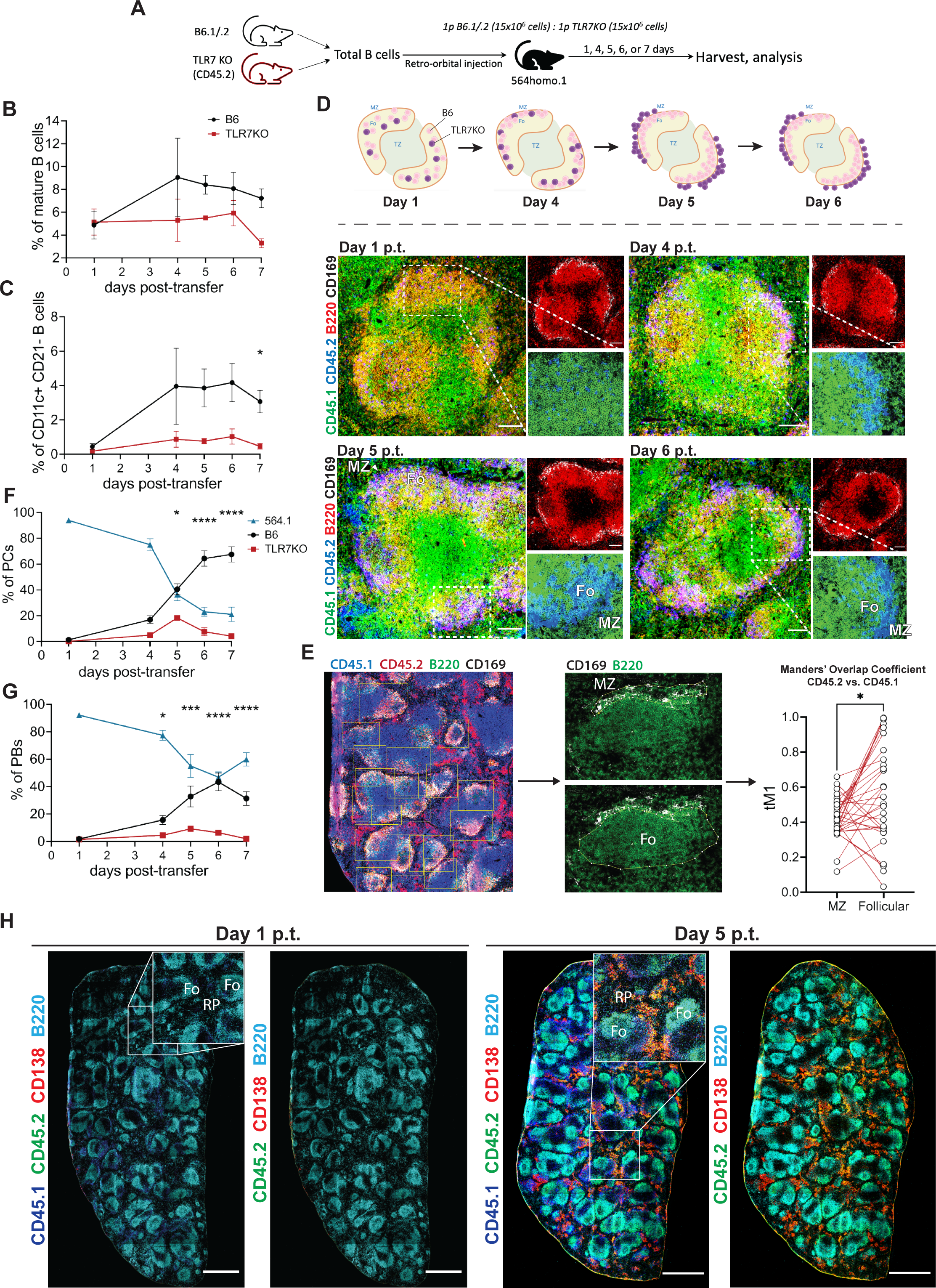
WT and *TLR7*-KO donor B cells occupy distinct splenic follicular niches and have divergent outcomes. (A) Experimental set up of competitive transfer time course. (B, C) Respective total mature and CD11c+ CD21- CD138- B cell time course plots from flow cytometry analysis at 1- (n=2), 4- (n=3), 5- (n=4), 6- (n=3), 7- (n=8) days post-transfer from at least 2 independent cohorts. Statistical analyses with Šidák’s multiple comparisons test. (D) IF imaging of host splenic white pulp at various days post-transfer. B220 and CD169 composite images show CD169+ MZ borders around B cell follicles; insets show composite of CD45.1 and CD45.2 near MZ-follicular borders. Fo=follicle, MZ=marginal zone. Scale bars = 100 µm. (E) Workflow of CD45.1 (channel 2) and CD45.2 (channel 1) co-localization analysis (Fiji Coloc 2 plugin) in transfer host spleens and tM1 quantification of MZ and follicular space. Representative follicle selections and MZ/Follicular outlines demonstrated on IF images. Analysis completed on 32 white pulps from two spleens at 5 days post-transfer. Statistical analyses with paired t test. (F, G) Respective PC (CD138+ B220-) and PB (CD138+ B220+) time course plots from flow cytometry analysis over 7 days of transfer. Statistical analyses with Šidák’s multiple comparisons test. (H) IF images of whole spleens from 564Igi transfer recipients at one (left) and five (right) days post-transfer at 30x magnification. Scale bars = 1mm. *=p<0.05, ***=p<0.001; ****=p<0.0001.

In terms of plasma cells (CD138^+^B220^-^), the WT and *TLR7*-KO cells again showed a clear divergence (**Fig. 2F**). Rapid EF PC differentiation began at 4 d.p.t. for both donor B cells, with WT PCs quickly outcompeting *TLR7*-KO cells to occupy approximately 60% of the PC compartment. Interestingly, while WT donor PBs steadily increased from day 4 post-transfer, they never exceeded 50% of the PB pool and contracted at 7 d.p.t., indicating that EF PBs may be an exhaustible, transient ASC reserve (**Fig. 2G, s2D-E**). This is also reflected in rapid expansion of the total PC compartment, compared to relatively static PB compartment (**Fig. s2C**). Immunofluorescence (IF) imaging showed significant accrual of donor ASCs in the splenic red pulp at 5 d.p.t., with minimal donor/host spatial segregation (**Fig. 2H**). In summary, competitive transfer time course elucidated the rapid co-optation of EF PC compartment by WT donors and further supports that TLR7-positive and -negative B cells take different developmental courses.

### Donor B cells exhibit a fixed division program and require at least 7 divisions for ASC differentiation

To assess donor B cell proliferation and their division profile, we labeled WT and *TLR7*-KO total B cells with Cell Trace Violet (CTV) and co-transferred at a 1:1 ratio into 564.1 recipients (**Fig. 3A**). Spleens were harvested at 5 d.p.t. for flow cytometry analysis and IF imaging to capture the peak EF differentiation window. We found that most donor B cells remained undivided in the B cell follicles, but some WT and *TLR7*-KO B cells went through at least 7 divisions after 5 days in the host (**Fig. 3B; s3A, B**). A smaller fraction of *TLR7*-KO B cells reached 7+ divisions relative to WT B cells, which is consistent with their reduced ASC differentiation (**Fig. 3B**). In fact, more *TLR7*-KO donor B cells were lost in competition to WT at each division, indicating that fewer *TLR7*-KO B cells were signaled to divide in each cell cycle (**Fig. 3C**). In agreement with recent findings by Scharer *et al*., the majority of ASCs (CD138^hi^) developed after the donor cells had undergone at least 7 divisions^40^ (**Fig. 3D**). Moderate upregulation of ASC- associated transcription factors BLIMP1 and IRF4 was also observed in late divisions, supporting ASC fate commitment of donor B cells (**Fig. 3E, F**)^41,42^. These results suggest that autoreactive ASC differentiation has a pre-determined division program that resembles foreign antigen exposure, and persistent TLR7 signaling licences EF B cell advancement through proliferation and toward ASC differentiation.

**Figure 3.**
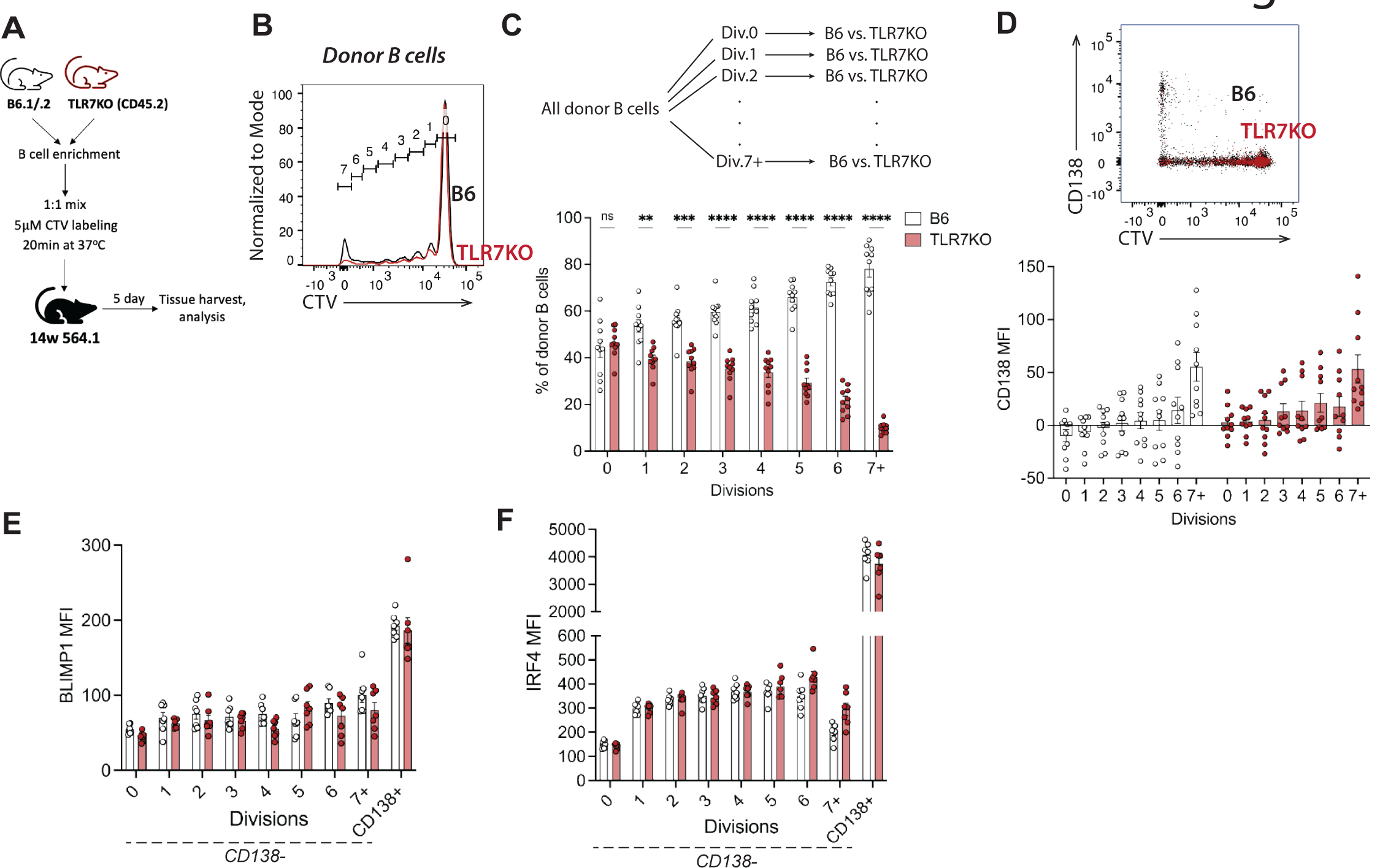
**At 5 days post-transfer, donor B cells exhibit a fixed division program and require at least 7 divisions for ASC differentiation.** (A) Experimental set up of donor B cell labeling with Cell Trace Violet (CTV) and competitive transfer. (B) Representative cytometry plot showing donor B6 and TLR7KO CTV histograms at 5 days post-transfer. (C) Gating strategy flow chart and frequencies of donor WT and TLR7KO B cells at every division (n=10). Statistical analysis with two-way ANOVA and multiple comparisons. (D) Representative flow cytometry plot of CD138 versus CTV and CD138 MFI at every division for B6 and TLR7KO donor cells. (E, F) BLIMP1 and IRF4 MFI of WT and TLR7KO donor B cells at every division (n=7). Statistical analysis with two-way ANOVA and multiple comparisons. *=p<0.05, **=p<0.01, ***=p<0.001, ****=p<0.0001.

### CD21^lo^ B cells exhibit rapid proliferation and demonstrate increased sensitivity to TLR7 deficiency

Lau *et al.* proposed that low CD21 expression represents a developmentally distinct stage for B cell differentiation and marks B cells that are recent GC emigrants primed for long-lived plasma cell differentiation in a vaccine response^43^. In SLE, both DN2 cells and ABCs express low level of CD21. Here, we found that both B6 and *TLR7*-KO donor B cells downregulated CD21 as they proliferated in the host and that *TLR7* KO did not affect donor B cell surface CD21 expression (**Fig. 4A**). For further analysis, we loosely divided the donor B cell compartment into CD21^+^CD23^-^ (MZ-like), CD21^mid^CD23^+^ (follicular), and CD21^lo^CD23^-^ (DN2-like, given high levels of CD11c, **Fig. 4B, s4D**)^44^, as well as into “resting” (division 0), “actively proliferating” (divisions 1-6), and “terminally divided” (divisions 7+). Strikingly, most MZ-like and follicular B cells were resting or actively proliferating, while cells that were terminally divided dominated the CD21^lo^CD23^-^ subset (**Fig. 4B, s4A**). Within the CD21^lo^CD23^-^ subset, we observed significantly more undivided *TLR7*-KO B cells than WT B cells, which instead dominated the divisions 7+ fraction, indicating that *TLR7* KO stalled donor cells at the resting stage (**Fig. 4C, D**). No significant difference was observed for MZ-like or follicular B cells between the two groups (**Fig. s4B, C**).

**Figure 4.**
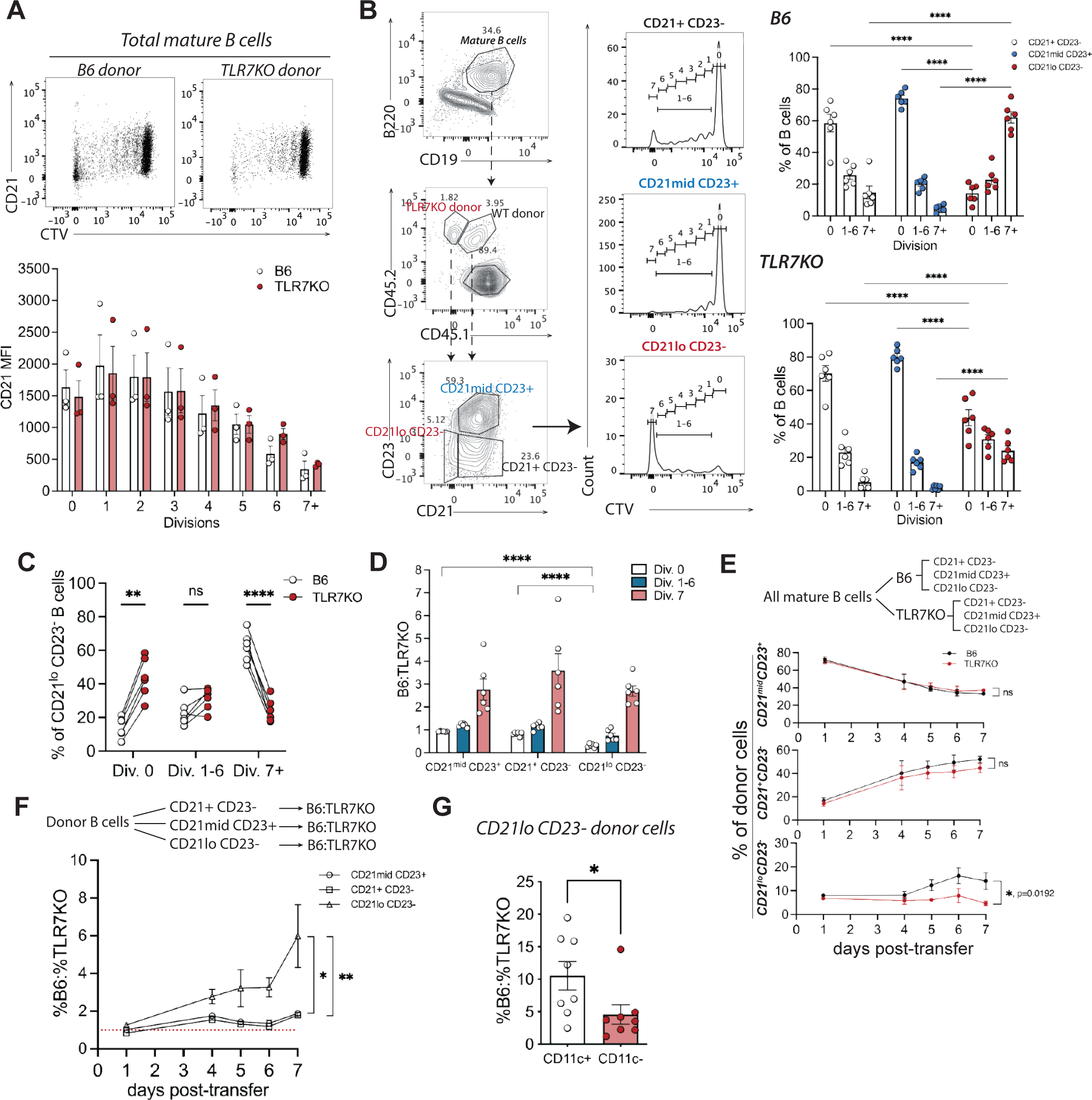
**CD21lo B cells exhibit rapid proliferation and demonstrate increased sensitivity to TLR7 deficiency.** (A) Representative flow cytometry plots of CTV versus CD21 for B6 and TLR7KO donor cells at 5 d.p.t. to 564Igi and corresponding quantification of CD21 MFI at each division (n=3). (B) Representative flow cytometry plots showing CTV histograms of CD21lo CD23-, CD21mid CD23+, and CD21+ CD23- donor subsets, and corresponding quantification of B6 and TLR7KO subset frequencies at 0, 1-6, and 7+ divisions (n=6). (C) Frequencies of B6 and TLR7KO donor CD21lo CD23- B cells with 0, 1-6, and 7+ divisions at 5 d.p.t. (D) B6-to-TLR7KO ratio of CD21lo CD23-, CD21mid CD23+, and CD21+ CD23- donor subset frequencies with 0, 1-6, and 7+ divisions at 5 d.p.t. (B, C, D) statistical analyses with paired t test. (E) Time course plot showing frequency changes of CD21lo CD23-, CD21mid CD23+, and CD21+ CD23- B6 and TLR7KO donor B cells over 7 days of transfer. Flow chart shows gating strategy used for quantification. (F) Time course plot showing changes in %B6 to %TLR7KO ratio within CD21lo CD23-, CD21mid CD23+, and CD21+ CD23- donor subsets over 7 days of transfer. (E, F) Area under the curve calculated for each subset between B6 and TLR7KO for Welch’s t-test analysis. ns=not significant. (G) %B6 to %TLR7KO ratio of CD11c+ and CD11c- CD21lo CD23- donor B cells at 7 days post-transfer. Statistical analysis with paired t *=p<0.05, **=p<0.01, ****=p<0.0001.

In fact, MZ-like and follicular B cells for both WT and *TLR7*-KO had similar proliferation kinetics over the course of 7-day transfer (**Fig. 4E**). However, WT CD21^lo^CD23^-^ B cell proportion increased from day 4 to 6 post-transfer, which synchronized with the peak time of donor ASC accumulation. This increase was absent in the *TLR7*-KO B cells, indicating that TLR7 specifically supports the expansion of CD21^lo^CD23^-^ B cells. Whereas the ratio of WT to *TLR7*-KO abundance persisted at below 2:1 throughout 7 days for MZ-like and follicular B cells, it was significantly higher for CD21^lo^CD23^-^ B cells especially at later timepoints, reflecting enhanced TLR7 sensitivity (**Fig. 4F**). Of note, when we split the CD21^lo^CD23^-^ group into CD11c+ (true DN2-like) and CD11c- subsets, then *TLR7* KO had a greater impact on the true DN2-like cells (**Fig. 4G**). Given their TLR7 sensitivity and proliferative tendency, we hypothesized that CD21^lo^CD23^-^ B cells were the precursor cells from which autoreactive EF ASCs differentiated and served as reservoirs of true DN2-like cells.

### CD21 downregulation during proliferation is reversible and can be restored by functional blockade of the receptor

Because CD21^lo^CD23^-^ B cells were predisposed toward EF ASC differentiation, we first hypothesized that CD21 was a negative regulator for early ASC differentiation. To test this, we adoptively transferred WT B6.1/.2 B cells mixed with B cells from a *CD21*-KO strain (mCD21^-/-^) into 564.1 mice and analyzed at 7 d.p.t. (**Fig. 5A**). Interestingly, CTV division profiling showed that mCD21^-/-^ B cells had a significant competitive advantage over WT in divisions 1 to 6, yet fewer mCD21^-/-^ B cells reached terminal divisions (7+) (**Fig. 5B**). The stalling during division coincides with CD21 acting in synergy with BCR to promote B cell activation and underscores the role of CD21 in regulating efficient B cell proliferation^31,45^. However, we found no difference between the frequencies of B6 and mCD21^-/-^ mature B cells and ASCs, indicating that constitutive *CD21* KO did not affect ASC production (**Fig. 5C, D**).

**Figure 5.**
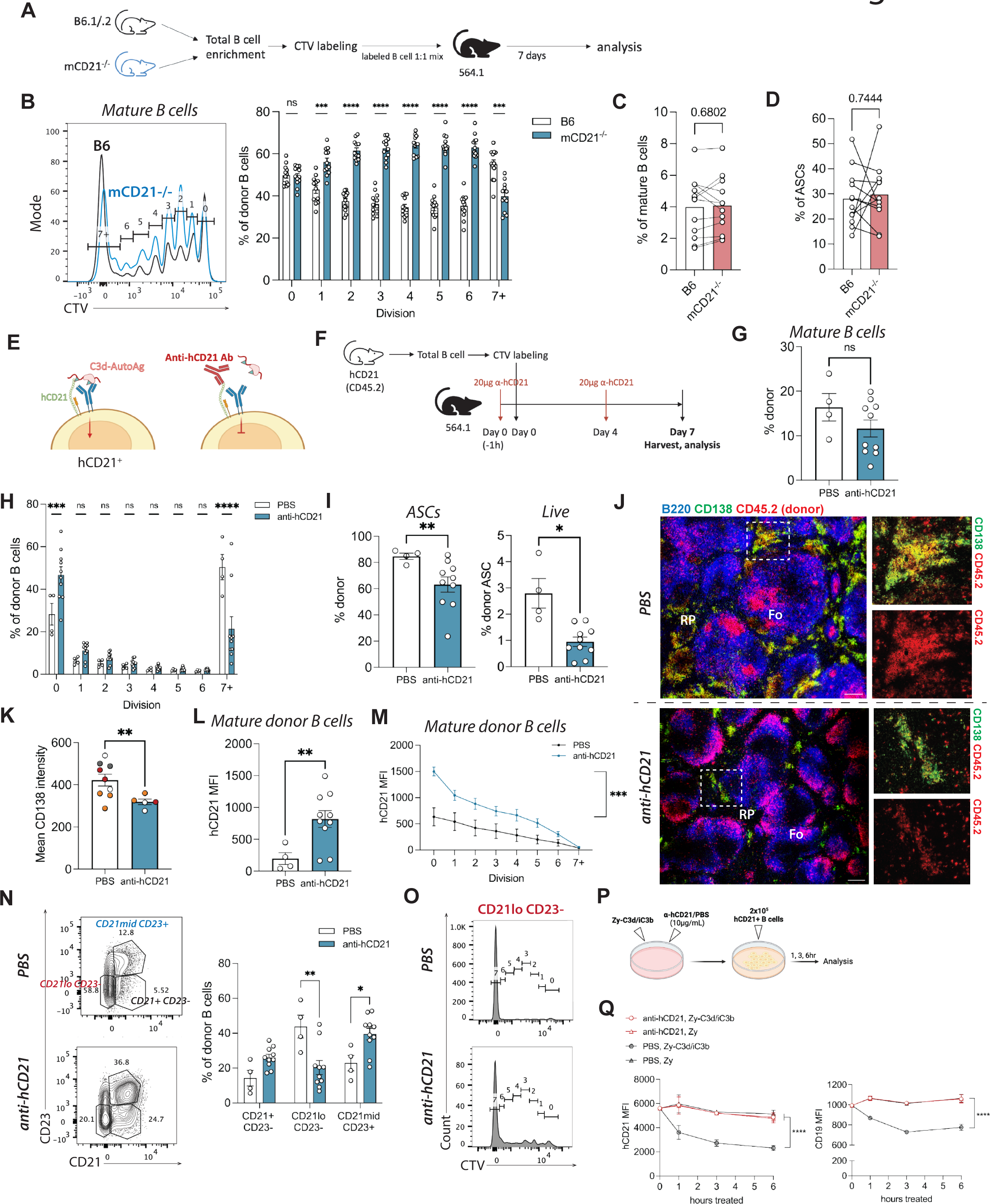
**CD21 downregulation during proliferation is reversible and can be restored by functional blockade of the receptor.** (A) Experimental set up of CTV-labeled WT and mCD21-/- B cell competitive transfer. (B) Representative CTV histograms of WT and mCD21-/- donor B cells and frequencies at every division. Statistical analysis with two-way ANOVA multiple comparisons. n=12 from at 3 independent cohorts. (C, D) Frequencies of WT and mCD21-/- mature B cells and ASCs (CD138+). Statistical analysis with paired t test and p values indicated on plots. (E) Diagram showing hypothesized mechanism of action of anti-hCD21 blocking antibody on target B cells. (F) Experimental set up of 7-day hCD21+ B cell blockade adoptive transfer. hCD21+ B cells were enriched, labeled with CTV, and transferred 564.1 recipients, which received 2 doses of blocking antibodies. (G) , (H), (I) Respective frequencies of total mature donor B cells (CD19+ B220+), donor B cells at each division, and donor ASCs as percent of total ASCs (left) or total live cells (right) with blocking antibody (n=10) or PBS (n=4) treatment from flow cytometry analysis of two independent cohorts. Statistical analysis with Welch’s t test (g, i) and two-way ANOVA multiple comparisons (h). ns = not significant. (J) Representative IF images of fresh frozen spleen cryosections from transfer recipient mice receiving PBS (top) or blocking antibody (bottom) at 30x magnification. Selected area of red pulp outlined with white box. RP = red pulp; Fo = follicle. Scale bars = 200 µm. (K) Mean CD138 intensity of spleen IF images from transfer recipient mice receiving PBS or blocking antibody per field of view. Each data point represents one field of view, and each unique color represents a different animal. (L, M) Respective hCD21 MFI quantification of total donor B cells and donor B cells at every division with blocking antibody or PBS treatment from flow cytometry analysis. Statistical analysis with area under the curve (l) and Welch’s t test (l, m). (N) Representative flow cytometry plots and relative frequencies of CD21lo CD23-, CD21+ CD23-, and CD21mid CD23+ donor B cells receiving blocking antibody or PBS treatment. Statistical analysis with Welch’s t test. (O) Representative CTV histograms of CD21lo CD23- B cells treated with blocking antibody or PBS. (P) Experimental schematic of *in vitro* hCD21 blockade assay. (Q) B cell surface hCD21 and CD19 MFI plots showing naive hCD21+ B cells treated with anti-hCD21 blocking antibody or PBS for 0, 1, 3, 6 hr in presence of Zymosan (Zy) or Zy-iC3b/C3d particles. Statistical analysis with area under the curve and Welch’s t test. *=p<0.05, **=p<0.01, ***=p<0.001; ****=p<0.0001.

Interestingly, we found that naïve mCD21^-/-^ mice had fewer B cells with anergic phenotypes and fewer MZ-like B cells, indicating that intrinsic differences in the B cell compartment of the two strains could complicate CD21 functional characterization (**Fig. s5A**). Hence, we next used a transgenic strain that expresses the human CD21 receptor on the mCD21^-/-^ background (hCD21^+^)^46,47^ and blocked its function with an antibody. The hCD21 blocking antibody targets the C3d binding site on hCD21, but not mCD21, and competitively inhibits the receptor’s interaction with complement-opsonized autoantigens, hence allowing donor-specific blockade (**Fig. 5E**). The recipient 564.1 mice were treated with two doses of 20 μg of monoclonal anti-hCD21 blocking antibody or PBS at day 0 and day 4. Splenic B cells enriched from hCD21^+^ mice were CTV labeled and adoptively transferred one hour after the first antibody treatment (**Fig. 5F**). Overall, we found no significant difference in mature B cell maintenance in the host, suggesting that CD21 blockade had a limited impact on donor B cell survival (**Fig. 5G**). However, hCD21 blockade prevented cells from reaching terminal divisions, increased the undivided fraction, and reduced donor-derived ASCs, indicating that restricting CD21 ligand engagement stalls donor B cell proliferation and compromises early ASC production (**Fig. 5H, I**). IF imaging of the host spleens also showed significantly fewer donor-derived ASCs upon hCD21 blockade (**Fig. 5J, K**).

Strikingly, blocking antibodies increased donor B cell hCD21 expression immediately after treatment (**Fig. 5L, M**), causing a restoration of CD21^+^ populations and a corresponding reduction of CD21^lo^CD23^-^ B cells while retaining their proliferation efficiency (**Fig. 5N, O**). The blockade also downregulated surface CD11c, suggesting reduced accrual of B cells with true DN2-like phenotypes (**Fig. s5B**). Increased surface CD19 expression was observed with blockade, supporting CD21 and CD19 acting together in a co-receptor complex to regulate B cell activation (**Fig. s5C**). Impact on surface IgM expression on donor B cells by blockade was insignificant (**Fig. s5D**). To validate that CD21 ligand engagement was directly causing CD21 downregulation on a molecular level, splenic hCD21+ naïve B cells were purified and cultured with Zymosan (Zy) particles coated with WT serum derived C3d/iC3b (Zy-C3d/iC3b) as described (**Fig. 5P**)^48^. B cell intrinsic CD21 was downregulated within 1-6 hr post- treatment in presence of Zy-C3d/iC3b but maintained in presence of naked Zy particles, with which C3KO serum was co-incubated as control. Strikingly, we found that hCD21 engagement with its cognate ligand caused immediate receptor downregulation and B cell rosette formation, which is effectively averted by addition of hCD21 blocking antibody (**Fig. 5P, s5E**). CD19 followed a similar surface expression pattern, indicating coupled activity with CD21 (**Fig. 5P**). Together, these data indicate that B cell CD21 engagement by and crosslinking with C3d/iC3b ligands initiate donor cell proliferation and differentiation and induce rapid downregulation of the receptor. The emergence of CD21^lo^ B cells could serve as an indicator of pre-activated B cells along the EF differentiation pathway.

### Follicular B cell–derived CD21^lo^ B cells are immediate precursors to autoreactive EF ASCs

While we provided evidence for CD21^lo^ B cells as immediate precursors for autoreactive EF ASCs, their upstream B cell origin remains elusive. To investigate such origin, we first individually sorted for MZ (CD21^hi^CD23^-^CD1d^+^IgM^+^), follicular (CD21^mid^CD23^+^), and transitional T1 B cells (CD21^lo^CD23^-^CD93^+^) from naïve B6 mice and transferred them into separate 564Igi hosts (**Fig. 6A, D**). The MZB have been hypothesized as primary candidates for EF ASC differentiation due to their close positioning and rigorous response to incoming blood-borne antigens^44^. However, they mostly retained their phenotypes in the host after 5.5 days, unlike the FoB (**Fig. 6B**). The MZB and follicular B cells produced similar numbers of ASCs (CD138^+^), but the abundance of FoB in the spleen suggests that the MZB may be a relatively smaller contributor to total ASC production (**Fig. 6C**, **s6A**).

**Figure 6.**
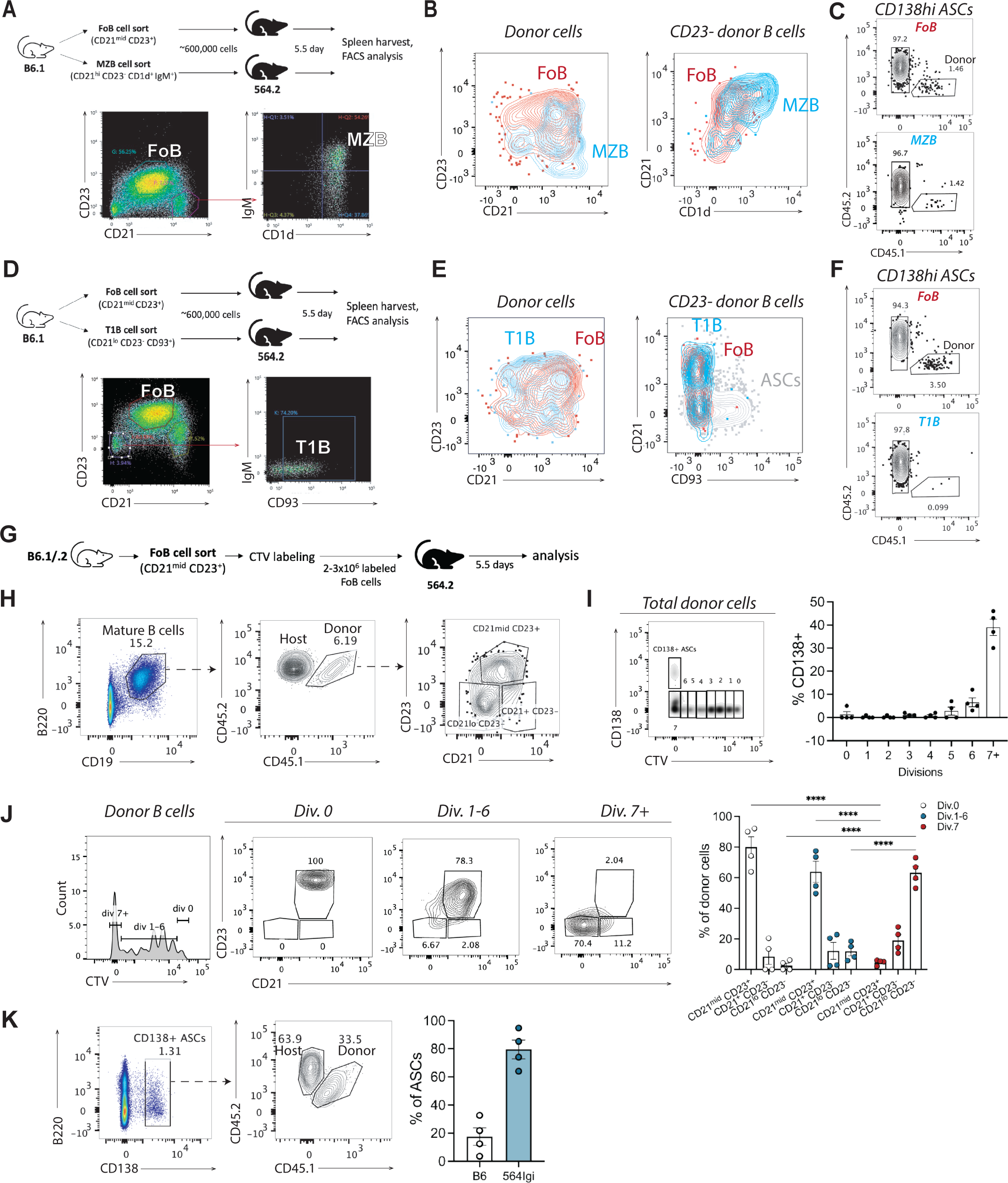
**Follicular B cell derived CD21lo B cells are immediate precursors to autoreactive EF ASCs** (A) Experimental set up and gating strategy of MZB / follicular B cell (FoB) transfers to 564.2 recipients. (B) Representative flow cytometry plots of donor B cells following FoB (n=5) and MZB (n=4) transfers showing follicular (CD21 versus CD23) and MZ (CD21 versus CD1d) B cell phenotypes. Populations sorted and transferred are indicated on respective plots. (C) Representative flow cytometry plots showing donor cell frequencies within ASC compartment from MZB / FoB cell transfers. (D) Experimental set up and gating strategy of T1B / FoB transfers to 564.2 recipients. (E) Representative flow cytometry plots of donor B cells following T1B (n=2) and FoB (n=3) transfers showing FoB (CD21 versus CD23) and T1B (CD21 versus CD93) phenotypes. Populations sorted and transferred are indicated on respective plots. (F) Representative flow cytometry plots showing frequencies of donor within ASC compartment from T1B / FoB transfers. (G) Experimental set up showing FoB sort, labeling and transfer to 564.2 recipients. (H) Representative flow cytometry plots showing frequencies of CD21lo CD23-, CD21mid CD23+, and CD23+ CD23- donor B cell subsets in 564.2 recipients following FoB transfer (n=4). (I) Representative flow cytometry plots of CTV versus CD138 and quantification of FoB derived CD138+ cells within each division. Statistical analysis with two-way ANOVA multiple comparisons. ****=p<0.0001. (J) Representative flow cytometry plots and quantification of donor B cell subset frequencies post FoB transfer at divisions 0, 1-6, or 7+. (K) Representative flow cytometry plots and quantification of donor and host ASC frequencies post FoB transfer.

Although the T1 B cells have partly overlapping surface phenotypes with CD21^lo^CD23^-^ B cells, they failed to differentiate into an appreciable number of ASCs (**Fig. 6F, s6B**)^49^. Instead, the 600,000 sorted T1 B cells lost CD93 expression, indicating maturation, and adopted mature follicular-like phenotypes at 5.5 d.p.t. (**Fig. 6D, E**). Consistent with these results, WT and *TLR7*-KO CD138^-^CD21^lo^CD23^-^ cells from total B cell transfers were mostly CD93^-^ (**Fig. s6C**). Nevertheless, they expressed surface CD24 at a level higher than the follicular B cells (CD21^mid^CD23^+^), indicating they may be more metabolically active^50^ (**Fig. s6D**).

Interestingly, at 5.5 d.p.t., sorted follicular B cells produced subsets resembling total B cell transfer, divided at least 7 times, and expressed CD138 mostly at 7+ divisions (**Fig. 6G-I**). Strikingly, while most dividing donor cells retained the follicular phenotype (CD21^mid^CD23^+^), most of the terminally divided cells were CD21^lo^CD23^-^ (**Fig. 6J**). Two to three million follicular B cells gave rise to an average of 20% of the total ASC compartment within 5.5 days (**Fig. 6K**). Together, these results support follicular B cells as a prominent contributor to early autoreactive ASCs while passaging through the intermediate state of CD21^lo^CD23^-^.

### B cell repertoire sequencing reveals developmental trajectories of follicular B cells toward autoreactivity

An important strength of the adoptive transfer model is its ability to support native repertoire diversity of the donor B cells. To track clonal evolution of autoreactive B cells along extrafollicular development, we adoptively transferred WT follicular B cells and performed bulk BCR heavy and light chain sequencing on the progeny donor CD21^lo^CD23^-^CD138^-^ (CD21^lo^) cells, follicular B cells, ASCs, and GC B cells at 6.5 days post-transfer (**Fig. 7A, s7A-B**). A fraction of the input cells was sequenced to establish the baseline clonal repertoire. Among the heavy chain sequences, the day 0 input follicular B cells exhibited the greatest diversity across different orders on the Hill diversity curve and the lowest clonal enrichment, followed by day 6.5 post-transfer follicular B cells (**Fig. 7B, s7C**). In contrast, CD21^lo^ cells demonstrated the lowest repertoire diversity and greatest clonal expansion, fitting their proposed role as effector cell precursors. Interestingly, ASCs showed greater clonal richness than GC B cells (q=0) with non-overlapping 95% CI, but lower overall repertoire diversity (q=1 and q=2), possibly due to somatic hypermutation (SHM) of the GC B cells^51^ (**Fig. 7B**).

**Figure 7.**
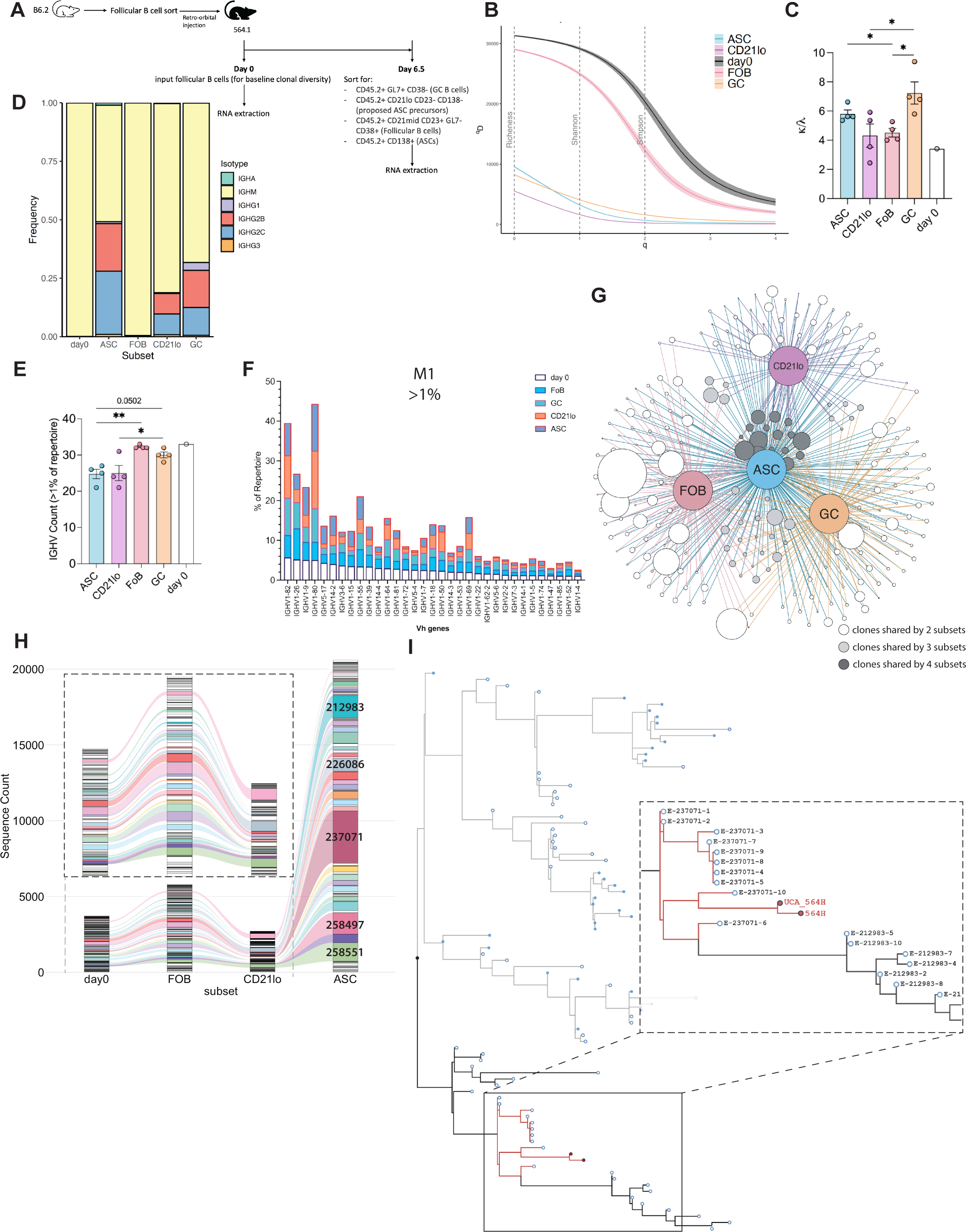
**B cell repertoire sequencing reveals developmental trajectories of follicular B cells toward autoreactivity** (A) Experimental set up of follicular B cell sort and transfer timeline; sorted donor populations are indicated with respective surface markers. (B) Repertoire diversity curve with Hill’s diversity index (^q^D) versus diversity orders (q). Subsets are colored respectively, with 95% confidence interval (CI) for each subset represented as shadowed areas of lines. ASC = ASCs, CD21lo = CD21lo cells, day0 = day 0 follicular B cells, FOB = day 6.5 follicular B cells, GC = GC B cells. (C) κ to λ light chain ratio for each subset. Ratio calculated from total number of unique sequences expressing Igκ divided by those expressing Igλ. Statistical analysis with Welch’s t test. *=p<0.05. (D) Plot of Ig isotype frequencies for each donor subset from four mice combined. (E) Counts of unique IGHV families for each donor subset contributing to more than 1% of subset repertoire. Statistical analysis with Welch’s t test. *=p<0.05, **=p<0.01. p value indicated on plot. (F) Vh gene repertoire frequencies of selected IGHV families from mouse 1 (M1) making up more than 1% of the input day 0 repertoire; stacked columns represent donor cell subsets at 6.5 d.p.t. with respective colors. (G) Qgraph illustrating shared clones between follicular B cells, CD21lo cells, ASCs, and GC B cells at 6.5 days post-transfer. Grey-scale circles represent individual clones, and colored circles represent subsets. Size of circles (in greyscale) is proportional to clone sizes. Connectors represent sharing of clones between different subsets. (H) Alluvial plot illustrating changes in clone sizes between subsets. Clones are pre-selected to only those shared by all four subsets. Thickness of bars with distinct colors represents absolute sequence counts within the clone. 5 most expanded ASC clones are annotated with their corresponding clone IDs. Inset illustrates enlarged region of plot showing only day 0 and 6.5 follicular B cells, and CD21lo cells. (I) Phylogenetic tree showing evolutionary relationships between selected donor ASC heavy chain sequences from 5 most expanded clones and 5 most restricted clones. Blue filled circles: sequences from restricted clones; blue empty circles: sequences from expanded clones. Inset shows clade including 564Igi, and 564Igi UCA heavy chain sequences and ASC sequences from two most expanded clones (#237071 and #212983). E = expanded, R = restricted.

Receptor editing is a tolerance mechanism that edits the κ light chain of an autoreactive cell for a λ light chain to dilute BCR autoreactivity^52,53^. Under autoimmune conditions, the preservation and expansion of κ light chain-expressing B cells may reflect selection for weakly autoreactive clones in the peripheral B cell repertoire. We found that GC B cells had the greatest κ-to-λ ratio that is also most distant from day 0 follicular B cells, implying advanced repertoire selection. CD21^lo^ cells and day 6.5 follicular B cells have lower κ-to-λ ratio than both ASCs and GC B cells, hence placing the two subsets early in development with less stringent selection (**Fig. 7C**). Class switch recombination of donor subsets support similar developmental relationships, with a shift from IgM+ naïve B cell in day 0 and day 6.5 follicular B cells to switched effector cells in CD21^lo^, GC B cells, and ASCs (**Fig. 7D, s7E**). Together, these results indicate that CD21^lo^ cells are developmentally more advanced than follicular B cells but precedes ASCs (and likely GC B cells).

VH gene family usage is another measure of repertoire diversity. ASCs and CD21^lo^ cells had lowest unique VH gene counts as compared to GC B cells and follicular B cells from both time points, suggesting selection and constricted VH gene diversity (**Fig. 7E**). In fact, they had prominent expansion of common VH genes (those that contributed to more than 1% of day 0 starting VH repertoire), such as IGHV1-82 and IGHV1-80 (**Fig. 7F**, **s7F**). Expansion of rare VH genes (e.g., IGHV1-19) that contributed to less than 1% of the starting repertoire was also observed.

Heavy chain clonal clustering and network analysis revealed that all four donor-derived B cell compartments demonstrated significant sharing of clones with varying degrees of expansion, indicating that same clones could commit to GC and/or EF fates (**Fig. 7G**). Fig. 7H illustrates clonal selection, expansion, and contraction from day 0 follicular B cells based on absolute sequence counts. The most expanded clones among ASCs are annotated with their respective numerical clonal identifications (**Fig. 7H**). To dissect possible autoreactive features of donor clones, a maximum likelihood phylogenetic tree was constructed with 8-10 randomly selected unique heavy chain sequences from the five most expanded and five most restricted ASC clones, together with the published 564Igi heavy chain sequence and its inferred unmutated common ancestor (UCA)^4^ (**Fig. 7I**). Remarkably, although derived from different germline V genes, the most expanded ASC clones, #237071 and #212983, were found on the same clade with the 564Igi and 564Igi UCA heavy chains, implying clonal selection of WT donors for 564-like heavy chains (**Fig. 7I**). Of note, sequences from clone #237071 used IGHV1-72, which was reported as the most frequently used mouse germline for autoreactive, DNA-binding antibodies in Nojima cell culture^54^ (**Table s1**). Overall, these results reflect rapid selection and expansion of proto- autoreactive clones in environment-driven break of tolerance and demonstrate early chronological evolution of follicular B cells and their progeny subsets.

## Discussion

In this study, we found that TLR7 is critical for sustaining WT autoreactive B cell differentiation in the EF pathway. While *TLR7* KO moderately affected the retention and survival of all donor B cells, it was most detrimental to EF ASCs and DN2-like cells. These observations coincide with Jenks *et al.,* who reported enhanced TLR7 responsiveness by human DN2 cells toward activation and PC differentiation^8^. In addition, many 564Igi Id+ B cells are functionally anergic and less responsive to survival signals than WT B cells upon engaging nuclear antigens through TLR7^12^, which could explain the gradual dominance of WT donor ASCs starting at 5 d.p.t., despite only a single WT B cell infusion.

The contrasting developmental fates of WT and *TLR7*-KO B cells could be attributed to their distinct placement in the white pulp and ability to uphold a fixed division program. Follicular exclusion and MZ migration of *TLR7*-KO B cells suggest that, while nuclear antigen sensing by TLR7 may not be essential for naïve B cells to survive, it does confer a competitive advantage for follicular access. This implies that passage through the B cell follicle promotes autoreactive B cell survival and differentiation even before GC development. Early immunization studies with T-independent antigen NP-Ficoll found that a substantial population of proliferating NP-specific B cells accumulated in the T cell zone, and a minor fraction also in the follicles^55^. Similar observations were made with AM14 Id+ MRL/lpr B cells^56^. While our model does not directly address the involvement of autoreactive T cells, we found that proliferating B cells localized to the follicles and not to the T/B border, indicating that prolonged T cell engagement may not be a requirement for autoreactive B cell proliferation and EF differentiation, perhaps due to pre-activation by surplus nuclear antigens from dead/dying cells. However, direct assessment of T/B engagement is needed, perhaps through the use of MHC-II deficient models.

We showed that naïve follicular B cells differentiate into substantial quantities of autoreactive ASCs within days, and that follicular B cell-derived CD21^lo^CD23^-^ B cells are an intermediate precursor to EF ASCs, with CD21 being instrumental to their proliferation earlier in the differentiation pathway. Moreover, proliferating donor B cells and ASCs downregulated CD21, consistent with the association between the absence of a functional CD21 and increased autoimmunity in murine SLE models that target nuclear antigens^57-5^^9^. As a B cell co-receptor, CD21’s engagement with CD19 enhances BCR- dependent activation by amplifying antigen receptor-mediated signal transduction^60^. We further showed that the loss of CD21 is reversible by blocking antigen engagement, which also stalled proliferation and effector cell development. The receptor’s elastic expression landscape could indicate a self-regulated receptor equilibrium. As surface CD21 expression decreases in proliferating donor B cells following antigen engagement, the loss of one BCR co-receptor may lead to increased dependence on TLR7 as the substitute co-stimulatory receptor, which is reflected in CD21^lo^CD23^-^ B cells’ increased sensitivity to TLR7 defect. Lau *et al.* showed that CD21^lo^ B cells upregulate negative regulators of BCR signaling, yet have no significant difference in the ability to flux intracellular calcium^43^. Strong TLR7 engagement by nuclear antigens may supply additional signals to counteract the inhibitory receptors and push CD21^lo^ B cells over the activation barrier. Future in-depth investigation into the connection between BCR signal strength, toll-like receptor dependence and ASC differentiation may generate new perspectives on activated B cell fate determination and antibody production. Finally, CD21 co- engagement with CD23 may imply coupled downregulation of the receptors on proliferating CD21^lo^CD23^-^ B cells, although we did not examine CD23 function in this study^61^.

MZ B cells are rapid generators of plasmablasts in response to T-independent blood-borne particulate antigens such as LPS and were thus attractive candidates for autoreactive EF ASC producers^62-6^^6^. However, follicular B cells being a main origin of donor-derived EF ASCs indicates that the proportional dominance of follicular B cells may override MZB’s rapid and robust response to autoantigens. Nevertheless, since MZB are a non-recirculating subset, the adoptive transfer system may hinder optimal donor MZB placement in the spleen^66,67^. Hence, while follicular B cells alone are sufficient to generate notable quantities of EF ASCs, they do not preclude MZB as distinct donors to the autoreactive ASC repertoire. On the other hand, while naive T1B did not differentiate into ASCs in days, they adopted follicular phenotypes and lost immature markers, indicating that they could develop mature naïve phenotypes in a foreign autoreactive host and potentially differentiate into ASCs in due time.

While we focused on TLR7 in this study, future studies should also explore other SLE-associated nuclear antigen sensors and complement receptors, such as TLR9, cGAS, and CD11c, as potential regulators of the EF response^68-7^^1^. In addition, while the adoptive transfer model provides a clean and modular system for the mechanistic study of GC-independent autoreactivity, we recognize that a single wave of donor B cells induces mild disease and limited tissue pathology. Chronic circulation of autoantigens in SLE enables continuous generation of ASCs and possible memory development, hence implying a potential for EF ASC-driven systemic autoreactivity. A critical next step would be to explore *in vivo* systems that reflect the complex outcomes of continuous EF ASC production and allow tracking of long-term disease progression, tissue pathology and clonal evolution.

In conclusion, our findings lend strong support for TLR7 to critically propagate efficient autoreactive EF ASC differentiation from naïve B cells. Our model further identifies CD21 as an antigen receptor that initiates autoreactive B cell proliferation before downregulation and resolves the apparent paradox of SLE-associated CD21^lo^ cell accrual. Together, the two receptors modulate tissue behavior of autoreactive B cells in break of tolerance, which underscore the complex innate-adaptive immunity interplay in rapid generation of autoantibodies. Furthermore, we demonstrate that B cells bearing potentially autoreactive BCRs can be rapidly selected in an autoreactive environment with minimal GC involvement. This study adopts new perspectives in examining SLE-associated disease phenotypes by focusing on the collaborative functions of B cell co-receptors. It may help elucidate repeated observations under varied settings and carry broader implications beyond autoimmunity, particularly in instances of infection where similar B cell phenotypes have been documented^72-7^^5^. With existing challenges in using B cell depletion therapies as a treatment for SLE, the use of hCD21-targeted blocking antibody offers practical implications as a potential inverse agonist to reversibly silence autoimmune B cells^76,77^. With accumulating evidence demonstrating the clinical significance of extrafollicularly-derived autoantibodies, we hope to advance the fundamental molecular understanding of the EF pathway in autoreactive B cell differentiation and to inspire development of target-specific therapies that enable selective modulation of immune regulators in place of universal suppression of immune cell function in the treatment of SLE.

## Methods

### KEY RESOURCES TABLE

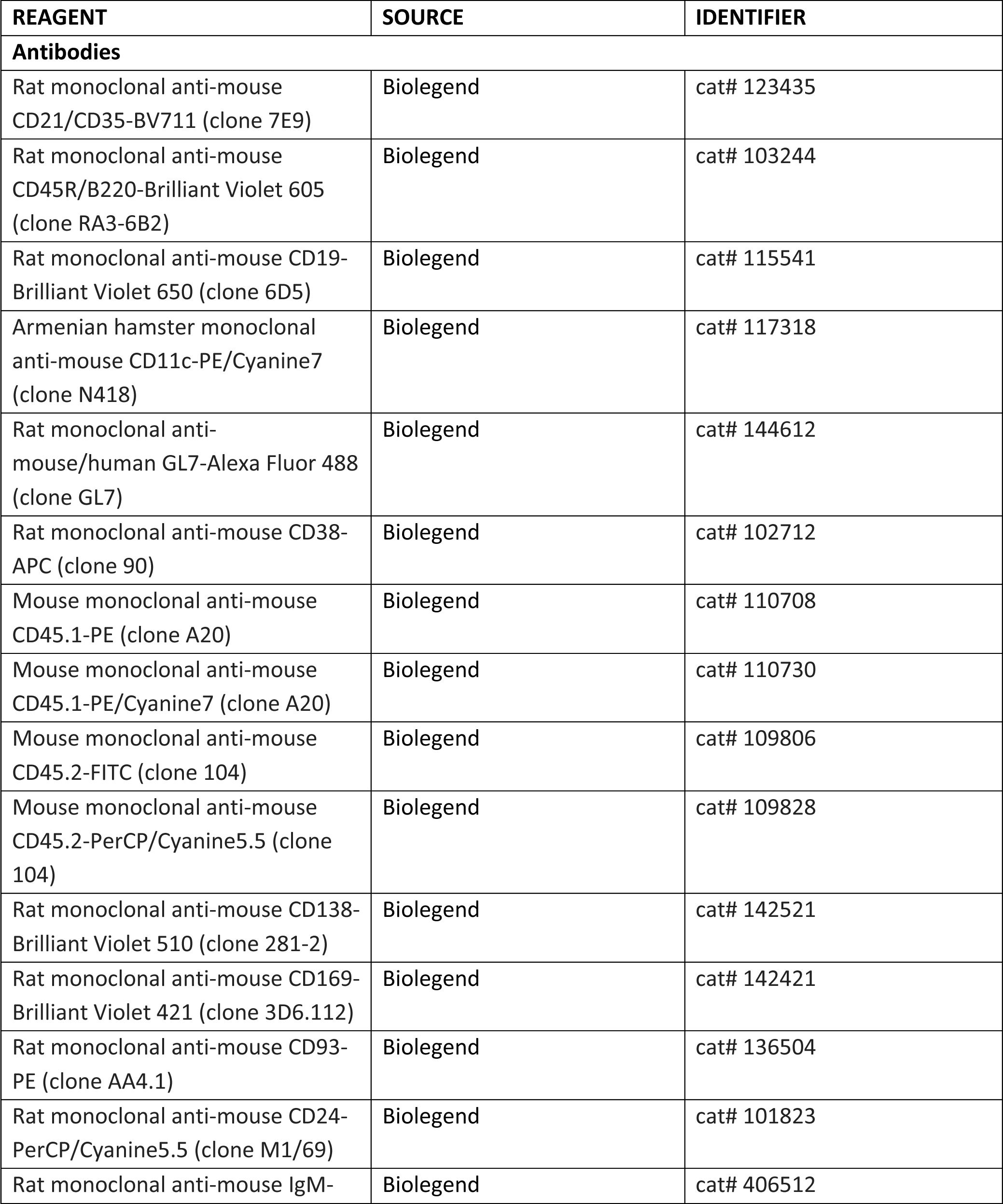
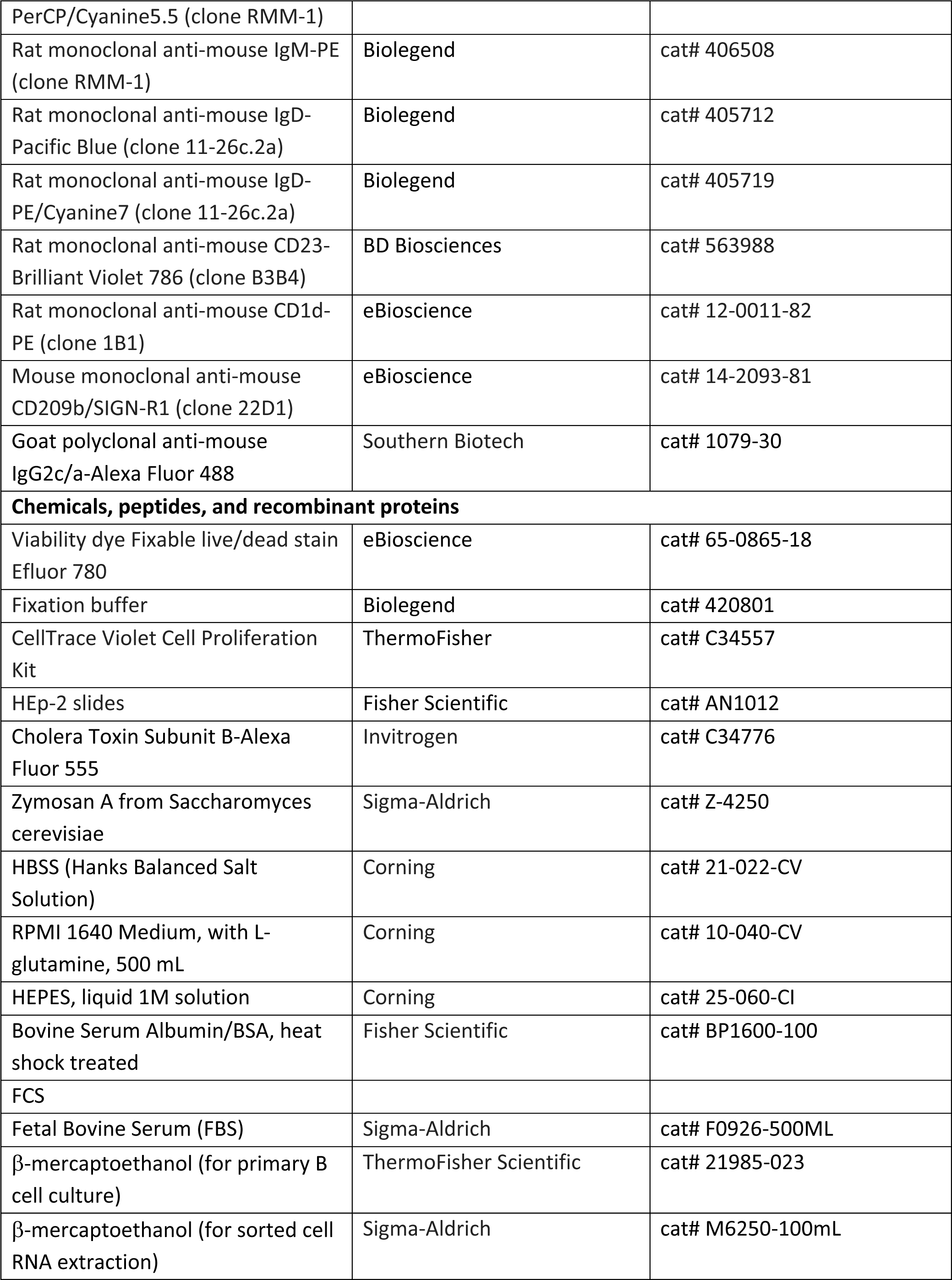
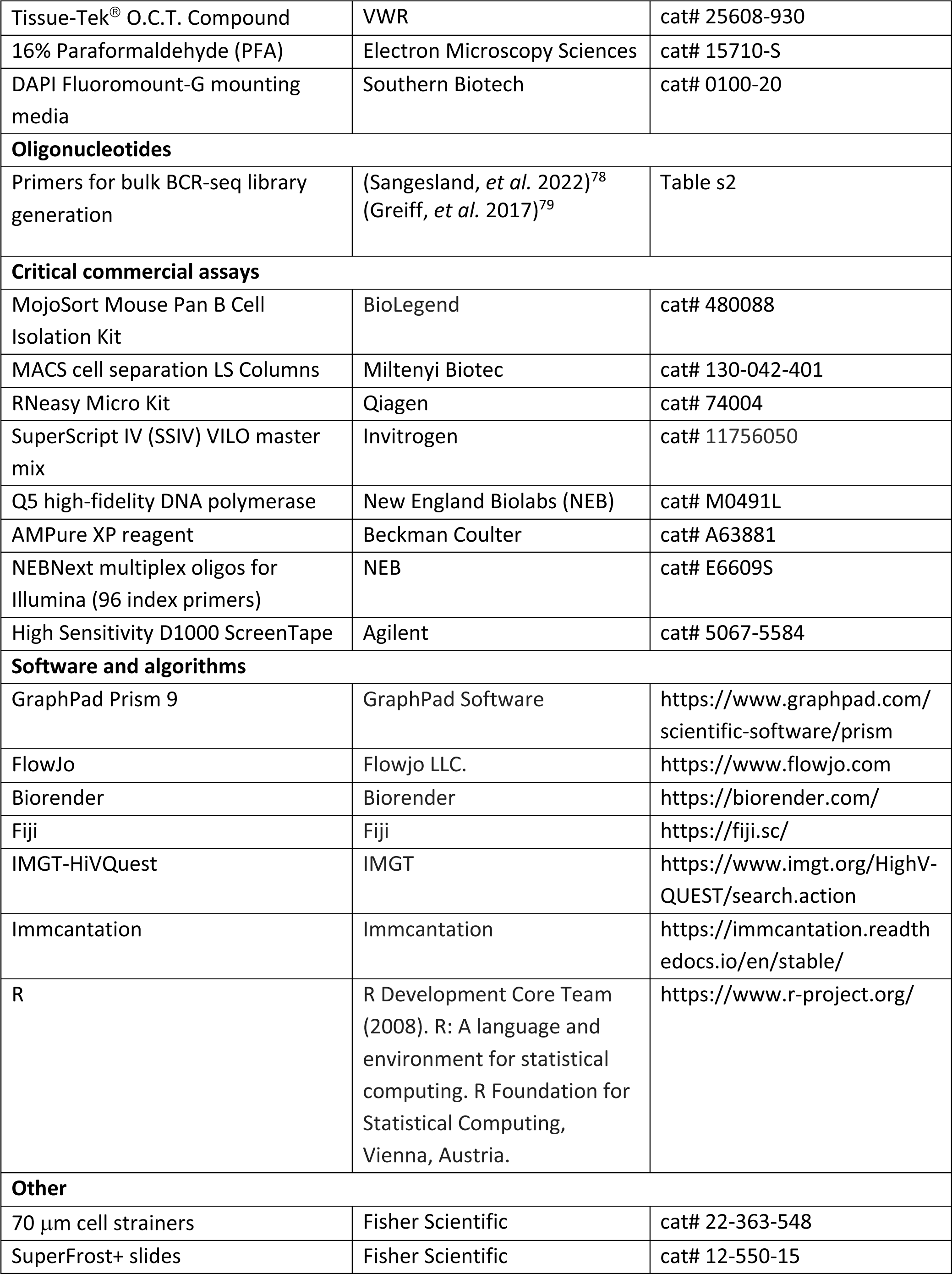
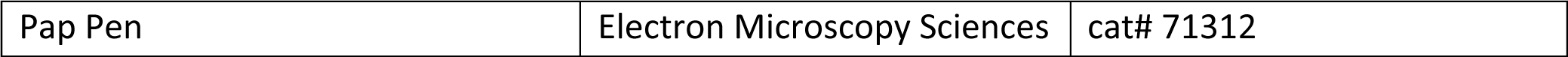

### EXPERIMENTAL MODEL AND SUBJECT DETAILS

#### Mice

C57BL/6J, B6.SJL (CD45.1), and TLR7 knockouts (B6.129S1-Tlr7tm1Flv/J) were obtained from Jackson Laboratories. mCD21-/- mice were generated and maintained in house^46,47^. The hCD21^+^ transgenic mice, gifted by Dr. Michael Holers (University of Colorado Denver) and maintained in-house, harbored hCD21 on a bacteria artificial chromosome and were bred onto the *Cr2-/-* background^47,80^. 564Igi mice^12^ were originally provided by Dr. Theresa Imanishi-Kari (Tufts University) and were maintained in-house. Mice were specific pathogen-free (SPF) and maintained under a 12-hr light/dark cycle with a standard chow diet. Both male and female mice were used. All animal experiments were conducted in accordance with the guidelines of the Laboratory Animal Center of the National Institutes of Health. The Institutional Animal Care and Use Committee of Harvard Medical School approved all animal protocols (protocol number IS00000111).

#### Antibodies and reagents

The monoclonal hCD21 blocking antibody (clone 658) was generated by Pfizer in a joint project with Boston Children’s Hospital (BCH) and shown to block human and mouse C3d binding to CD21 at sub- nanomolar affinity in human and cynomolgus monkey.

#### Flow Cytometry

Spleens and LNs were harvested into ice cold MACS buffer (0.5% bovine serum albumin/BSA, 2mM EDTA in 1x PBS) and mechanically dissociated using pestles in 1.5 mL Eppendorf tubes. Samples were then filtered through 70 μm cell strainers (Fisher Scientific) and centrifuged at 1400 rpm at 4°C for 5min. LN cell pellets were resuspended in MACS buffer. Spleen cell pellets were resuspended in RBC lysis buffer (155 mM NH4Cl, 10 mM NaHCO3, 0.1 mM EDTA), incubated for 5 min on ice and centrifuged as before. Splenocyte pellets were washed once with MACS buffer, centrifuged and resuspended in MACS buffer. Single-cell suspensions were added to 96-well round-bottom plates, centrifuged and resuspended in 50 μL antibody staining mix (1/200-1/400 diluted fluorophore-conjugated/biotinylated Ab and 1/1000 diluted viability dye in MACS buffer). Samples were incubated for 30 min at 4°C, followed by addition of 150 μL of MACS buffer. Plates were subsequently centrifuged at 1400 rpm for 5 min, and the supernatant was flicked out of the plates. For the two-step staining procedure, streptavidin- conjugated secondary antibody mixture was added and incubated for 30 min at 4°C. The samples were then fixed for 20 min at room temperature in the dark with 100 μL of fixation buffer (BioLegend). For surface staining only, 100 μL of MACS buffer was added to each well and centrifuged 1400 rpm for 5 min. The supernatant was removed, and cell pellets were resuspended with 150 μL of MACS buffer ready for acquisition. For intracellular staining, fixed samples after surface staining were then permeabilized with 200 μL of 10x Intracellular Staining Perm Wash Buffer (diluted to 1X in dH2O) for 15 min at room temperature. 1x Perm Wash Buffer was then removed after centrifugation at 1400 rpm for 5 min. Permeabilized cells were resuspended in 50 μL of intracellular antibody mixture in Perm Wash Buffer. Samples were washed by addition of 150 μL of 1x Perm Wash Buffer and centrifugation. After the supernatant was removed, samples were resuspended with 150 μL of MACS buffer and kept at 4°C until ready for acquisition. Sample acquisition was performed on one of three instruments, depending on the experiment: a 3-Laser 8-color Canto-SORP analyzer with HTS equipped with 488nm, 405nm, and 588nm lasers, a 4-Laser 17-color LSR II analyzer with HTS equipped with 405 nm, 355nm, 594nm, and 488nm lasers, and a 5-Laser 17-color LSR II SORP analyzer with HTS equipped with 355nm, 405nm, 488nm, 561nm and 640nm lasers. Flow cytometry data analyses were performed with FlowJo software.

#### B cell adoptive transfer

Donor B cells were enriched using MojoSort Mouse Pan B Cell Isolation Kit according to manufacturer protocols. Briefly, total splenocyte single cell suspensions from whole spleen were prepared as described and equilibrated to 100x10^6^ cells/mL in MACS buffer followed by 15 min incubation with biotinylated antibody cocktail and 15 min incubation with streptavidin magnetic beads, both on ice. The cells were then passed through the MACS cell separation LS Columns (Miltenyi Biotec) fixed on magnet and collected into MACS buffer. Cells were then centrifuged, resuspended in 0.5 to 1 mL of MACS buffer and counted. The desired number of cells were retrieved, pelleted, and resuspended in sterile HBSS buffer with 10 mM HEPES, 1mM EDTA, 2% FBS, ready for transfer. Adoptive transfer was performed by single-eye retro-orbital injection at 100 μL volume with 0.5 mL Lo-Dose Insulin Syringes with 29 G needle (Becton Dickinson).

#### Cell trace staining for *in vivo* proliferation analysis

Cell trace violet/CTV staining of B cells was completed per manufacturer recommended conditions (Thermo Fisher). Enriched splenic B cells (method as above) from donors were incubated in 5-20 μL of 5 μM CTV dye working solution. CTV working solution was prepared by adding 20 μL DMSO to CTV stock vial to generate 5 mM stock solution and transferring 10 μL of stock solution to 10 mL of PBS in a 15 mL conical tube. The working solution was mixed thoroughly by inverting the tube, and the cell pellet was first carefully and thoroughly resuspended in PBS, then added to the CTV solution. The cells were then incubated in a 37°C water bath for 10 min (tube inverted every ∼5 min), placed on ice, and transferred to a 50 mL conical tube. 3-4 volumes of RPMI+10% FBS was added to cells to quench the remaining dye. Cells were centrifuged at 1400 rpm for 5min at 4°C, and cell pellets were resuspended in 500 μL - 1 mL RPMI media + 10% FBS. Cells were counted; the desired number of cells were retrieved and transferred to a new, sterile Eppendorf tube, pelleted, and resuspended in HBSS buffer with 10 mM HEPES, 1mM EDTA, 2% FBS for 100 μL per injection, ready for transfer.

#### FACS sorting

Spleen cells for sorting were prepared from tissue and surface-stained same way as in flow cytometry analysis. For follicular B cell sorting, total splenic B cells were first enriched prior to sorting. Single-cell suspensions, FMO controls, and single-color stain compensation samples were resuspended in sterile HBSS buffer with 10 mM HEPES, 1 mM EDTA, and 2% FBS at appropriate concentrations and filtered immediately prior to sorting. Cells were sorted into RPMI media with 10% FBS and kept on ice until use. Two sorters were utilized: 5-laser (488nm, 406nm, 592nm, 640nm, and 355nm) FACSAria-SORP cell sorter with PMT-forward scatter option; Sony cell sorter SH-800Z with 4 laser (405nm, 488nm, 561nm, and 633nm), 6-color configuration and 70 μM sorting chips.

#### Immunofluorescence confocal microscopy and image analysis

Fresh tissue was embedded in Tissue-Tek^®^ O.C.T. Compound (VWR) after harvest, and rapidly frozen on dry ice. Tissue blocks were stored at -80°C until ready for cryosection. 14 μm tissue cryosections were mounted on SuperFrost+ slides (Fisher Scientific). For immunofluorescence staining with surface and intracellular markers, slides were dried at room temperature for 20 min and each section on the slide was outlined with hydrophobic barrier with Pap Pen (Electron Microscopy Sciences) and fixed with 4% PFA at RT for 10min. After rinsing with PBS, slides were then permeabilized with 0.2% Triton in 1x PBS for 3-5 min at RT, washed 3 times with PBS, 0.05% Tween-20 (PBST) and incubated for 1 hour in blocking buffer (3% BSA, 5% fetal calf serum/FCS in 1x PBS). This was followed by incubation with primary Ab mixture in blocking buffer overnight at 4°C. For the two-step staining procedure, fluorescently conjugated secondary Ab mixture in blocking buffer were added to slides for 1 hr incubation at room temperature. The slides were then washed 3 times with PBST, mounted in Fluoro- Gel (Electron Microscopy Sciences), applied with coverslips, and sealed with clear nail polish. Confocal Imaging of spleen sections was performed with Olympus FV3000R resonant scanning confocal microscope, which is equipped with 4 laser lines (405, 488, 514, and 633 nm) and ultra-sensitive GaAsP detectors with full spectral imaging. Images were acquired with 30x silicone oil objective and frames stitched with the Olympus CellSens software. Image processing and signal intensity measurements were completed with Fiji (ImageJ).

#### HEp-2 Immunofluorescence assay

12-well HEp-2 slides were removed from the pouch and equilibrated to room temperature for 5-10 min. Hydrophobic barriers were marked around each well with a PAP pen. Sample sera (from adoptive transfer or naïve mice) were diluted 1/20 or 1/50 in PBS, and 30 μL of diluted sera were applied to each well to form a “dome”. The slides were incubated at room temperature for 1 hr. Sera was removed by aspiration followed by 2x 5 min wash with blocking buffer (0.1% Tween-20, 0.5% BSA in 1x PBS). Avoid over-drying slides in the process. Slides were then incubated in secondary antibody mixture with 1/400 anti-IgG2c/a-AF488 and 1/400 Cholera Toxin B-AF555 in blocking buffer at room temperature for 1 hr. The antibodies were rinsed off with blocking buffer, followed by 2x 5 min washes. Without over-drying slides, the blocking buffer was completely removed. DAPI Fluoromount-G mounting media (Southern Biotech) was added to slides, coverslips were applied, and sealed with nail polish ready for imaging. Images were acquired with Olympus FV3000R resonant scanning confocal microscope with 100x silicone oil objective.

#### *In vitro* hCD21 blockade assay

Untouched resting/naive B cells were isolated from whole spleens of hCD21^+^ mice using the Miltenyi B cell isolation kit. Premix of anti-hCD21 blocking antibody (10 μg/mL) was prepared in RPMI media with 10% FBS and 1xβ-mercaptoethanol (complete RPMI). Zy-iC3b/C3d conjugation was performed as described with slight modification^48^. Serum samples were collected from WT and C3KO mice. 500-700 μL sera was incubated with Zymosan A beads at final concentration of 1mg/mL for 30 min at 37°C for full complement consumption and activation. The sera/Zy mixture was mixed by tube inversion every 10-15 min to prevent sedimentation. 0.5M EDTA was then added to the mix to final concentration of 10 mM and incubated at 37°C for 30 min to stop further reaction. The serum/Zy mixture was then centrifuged for 5 min at 1400 rpm. The supernatant was discarded, and Zy-iC3b/C3d pellet (or Zy with C3KO serum control) was resuspended in complete RPMI. 20 μL Zy-iC3b/C3d suspension was added to 96 well plates containing 80 μL blocking antibody premixes. 2x10^5^ naïve B cells were added to each well and incubated for 1, 3, and 6 hr before flow cytometry analysis.

### Bulk BCR sequencing

#### Follicular B cell transfer and B cell subset sorting

CD21^mid^CD23^+^ follicular B cells were sorted from enriched splenic B cells and polled from six B6 mice, resuspended in sterile HBSS buffer with 10 mM HEPES, 1mM EDTA, 2% FBS and transferred i.v. (retro- orbital) into four 564Igi recipients. Each recipient received ∼7.5x10^6^ cells. A fraction of sorted follicular B cells was saved for RNA extraction on the same day. At days 6.5 post-transfer, spleens of recipient mice were harvested, and donor B cell subsets (CD21^lo^, ASCs, GC B cells, and follicular B cells) were sorted from splenocyte single-cell suspensions into 5 mL FACS tubes. The sorting panels included CD21, CD23, CD138, CD45.1, CD45.2, CD19, B220, GL7, CD38, and Viability dye eFluor 780. Sorted cells were kept on ice in 0.5 mL RPMI media with 10% FBS until ready for RNA extraction. RNA extraction was completed the same day after sorting with RNeasy Micro Kit (Qiagen). For each sample, sorted cells were transferred to and centrifuged in 1.5 mL Eppendorf tubes at 1400 rpm for 5 min. Supernatant was carefully and thoroughly removed by aspiration, and 350 μL buffer RLT with 1% β-mercaptoethanol was added immediately to the cell pellet for lysis and pipetted thoroughly to mix. 350 μL freshly prepared, RNase-free 70% ethanol was added and mixed well by pipetting. The sample was immediately transferred to the RNeasy MinElute spin columns in 2 mL collection tubes and centrifuged for 30 s at 12,000 x g on a table-top centrifuge. Flow-through was discarded. 350 μL of Buffer RW1 was added to the spin column and centrifuged for 30 s at 12,000 x g. Flow-through was discarded. For DNA digestion, 10 μL of DNase I stock solution in 70 μL of Buffer RDD (gently mixed) was added directly to the spin column membrane and incubated at room temperature for 15 min. 350 μL of buffer RW1 was added to the spin column, followed by 30 s centrifugation at 12,000 x g. The column was placed in a new 2 mL collection tube, 500 μL of Buffer RPE was added, and centrifuged for 30 s at 12,000 x g. Flow-through was discarded. 500 μL of freshly prepared, RNase-free 80% ethanol was added to the spin column, followed by centrifugation for 2 min at 12,000 x g to wash. The spin column was placed in a new 2 mL collection tube and centrifuged at full speed for 5 min with lid open. Collection tube and flow-through was discarded. RNA was eluted with 14 μL RNase-free water into a new, RNase-free 1.5 mL collection tube with 1 min centrifugation at full speed. RNA was stored at -80°C until ready for library preparation. Day 0 follicular B cell RNA was quality controlled by TapeStation.

#### Library preparation

Isolated B cell RNA (14 μL) was reverse transcribed using SSIV VILO (Invitrogen), which contains both poly(dT) and random hexamers for priming, under the following conditions: 25°C for 10 min, 50°C for 20 min, 85°C for 5 min. The product of the reverse transcription reaction was purified with AMPure XP beads at a 1:1 bead to sample ratio (Beckman Coulter). Purified cDNA was split in half to amplify the heavy and light chains in separate reactions. The heavy chains were amplified in a first PCR step with 19 forward primers that anneal in the VH framework 1 region^81^ and a mixture of mouse IgG, IgM and IgA reverse primers that contain partial adapter sequences^78,79^ (**Table s2**). The light chains were first amplified using a mix of 3 forward primers that anneal in the Vk or Vλ framework 1 region^82^ and 2 reverse primers specific to Ck or Cλ (**Table s2**). The first PCR conditions were similar to those previously reported^79^, using Q5 polymerase (NEB): 98°C for 30s; 4 cycles of 98°C for 10s, 50°C for 20s, 72°C for 30s; 4 cycles of 98°C for 10s, 55°C for 20s, 72°C for 30s; 10 cycles of 98°C for 10s, 50°C for 20s, 72°C for 30s; 72°C for 2 min. The heavy and light chain PCR products were purified separately with AMPure XP beads at a 0.8:1 bead to sample ratio. The remainder of the adapter sequences were added to the heavy and light chains separately by a two-step PCR reaction with Q5 using the NEBNext index primers (NEB) 98°C for 30s; 12 cycles of 98°C for 10s, 72°C for 60s, 72°C for 5 min. The heavy and light chains from the same original sample contained the same index. The final PCR products were purified by AMPure XP beads at a 0.8:1 bead to sample ratio and quality controlled by TapeStation (D1000, Agilent, cat# 5067-5584). The samples were pooled at equimolar concentrations and further purified by gel extraction and AMPure XP bead clean up.

#### Sequence analysis

Pooled cDNA libraries were submitted to the Biopolymers Facility at Harvard Medical School for qPCR validation and sequenced with an Illumina MiSeq sequencer for 20 to 25 million reads, 600 cycle- medium kit length with paired-end sequencing. Raw, demultiplexed FASTQ sequence files were distributed and pre-processed on the HMS O2 Cluster. Paired-end reads were merged using the pRESTO pipeline under the Immcantation framework^83^. FastQC and MultiQC reports were generated for raw and merged reads. Primer masking, duplicate sequence removal, and repeated sequence filtering were performed with pRESTO. Processed sequences were submitted to IMGT/HighV-QUEST for alignment and annotation^84^. IMGT output was processed with Change-O to remove non-productive sequences. Heavy and light chain sequences were parsed into separate files with Change-O^85^. IgBLAST database was configured from IMGT reference sequences and used for germline reconstruction. The shazam R package was used for calculating the nearest neighbor distances and clonal clustering threshold^85^. Clonal clustering was completed with Change-O and a threshold of 0.045 was applied (**Fig. s7D**). The alakazam R package was used for the calculation and generation of clonal diversity and abundance curves^86^. Ig isotypes were obtained through MiXCR analysis pipeline for MiLaboratories Human IG RNA Multiplex Kit. Plot demonstrating sharing of clones between subsets was generated with the qgraph R package^87^. Clonal expansion alluvial plot was generated using the ggalluvial R package^88^. IGHV family frequencies were visualized using GraphPad Prism 9. The Phylogenetic tree for selected sequences from expanded and contracted clones was derived using CLUSTALW multiple sequence alignment tool and verified using the Abalign software^89^.

### Quantification and statistical analysis

All statistical analyses were performed in GraphPad Prism version 9, and specifically indicated throughout text and figure legends.

### Data and code availability

Raw sequence data for bulk BCR-sequencing have been deposited to Sequence Read Archive (SRA) with the accession number PRJNA1051189. Computer code for BCR repertoire analyses can be accessed at GitHub (https://github.com/Danni-Zhu/DZ_CD21lo_BCR-seq; https://github.com/yixiangD/sequencing/tree/main).

## Acknowledgements

We thank all members of the Carroll Laboratory for helpful discussion and feedback on the project and manuscript, and E. Carroll for administrative and managerial assistance. We thank V. Haridas, J. Moore (former director) and A. Agarwal from the PCMM Flow and Imaging Cytometry Resource, H. Leung from the Optical Microscopy Core, staff of the BCH/Cardiology Imaging and staff of the Harvard Biopolymers Facility for continuous technical assistance. We acknowledge D. Pascual and other staff members of the Warren Alpert Building and Harvard School of Public Health animal facilities for mouse colony maintenance. We are grateful to A. Marshak-Rothstein, J. Kagan for discussion, support, and advice regarding the project and critical reading of the manuscript, C. Usher for help editing the manuscript, C. Jensen and H. Meng from the Immcantation group for prompt advice on BCR sequence analysis, and A. Rattan for assistance with hCD21 blocking antibody preparation. We acknowledge Pfizer for generating the monoclonal blocking antibody against hCD21. This work is supported by NIH grant R01AR074105. D. Y. Zhu is supported by CIHR Doctoral Foreign Study Award 202110DFD-475148-94277. A. G. Schmidt is supported by NIH grants, R01AI146779 and P01AI089618. D. Lingwood is supported by NIH grants, R01AI137057, R01AI153098, R01AI155447.

## Author Contributions

D.Y.Z. and M.C.C. conceived the project. D.Y.Z., C.C., and M.C.C. conceptualized and planned most experiments. D.Y.Z. performed experiments and data analysis/interpretation with crucial assistance from D.P.M. for bulk BCR-seq methodology development, library generation, and data interpretation, and Y.D., F.A.N.M., and C.C. for BCR-seq data analysis and interpretation. M.M. performed genotyping of all mouse strains. M.C.C., C.C., D.L., and A.S. supervised the project and provided critical feedback.

D.Y.Z. wrote the manuscript. All authors provided feedback on manuscript.

---

**Figure S1.**
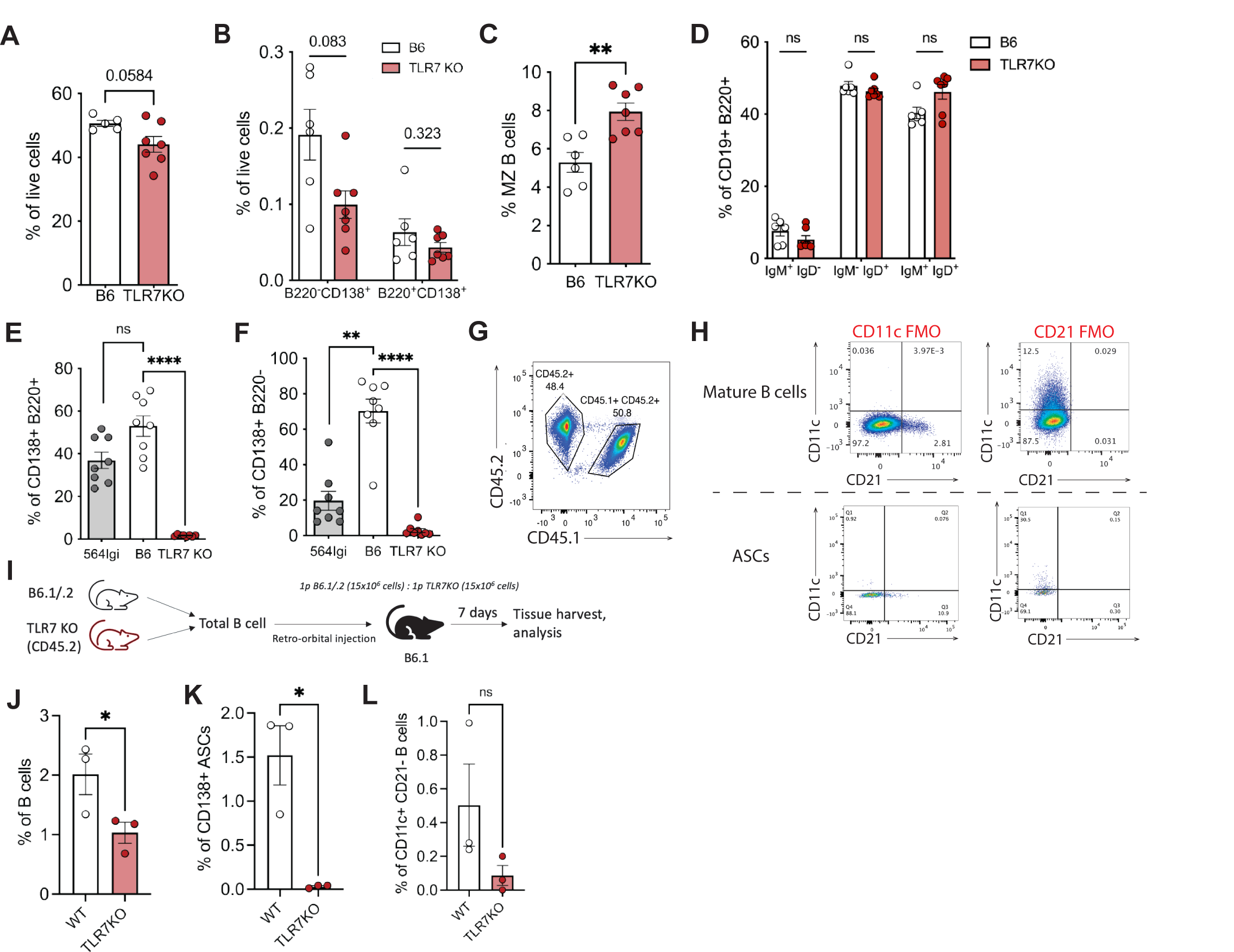
TLR7 drives efficient development of autoreactive B cells and EF ASCs with a DN2-like phenotype. related to Figure 1. (A, B, C) Splenic mature B cell, PBs (CD138+ B220+), PCs (CD138+ B220-), and marginal zone B cell frequencies of 14-20w old naive B6 (n=5) and TLR7KO (n=7) mice from flow cytometry analysis. (D) IgM+ IgD-, IgM- IgD+, and IgM+ IgD+ B cell frequencies of 14-20w old naive B6 and TLR7KO mice from flow cytometry analysis. (E, F) Respective frequencies of donor and host PBs and PCs at 7 days post-transfer (n=8) from flow cytometry analysis. (G) Representative flow cytometry plot showing input donor B cell mix with equal proportions of WT(CD45.1/.2) and TLR7KO (CD45.2) B cells. (H) Representative flow cytometry plots of CD11c and CD21 FMO controls used for CD11c+ CD21- B cell and ASC gating. (I) Experimental set-up of control competitive transfer to naive B6 recipients. (J, K, L) Respective frequencies of WT and TLR7KO donor mature B cells (CD19+ B220+), ASCs, and CD11c+ CD21- B cells in control competitive transfer (n=3). Statistical analysis with paired t test. ****=p<0.0001.

**Figure S2.**
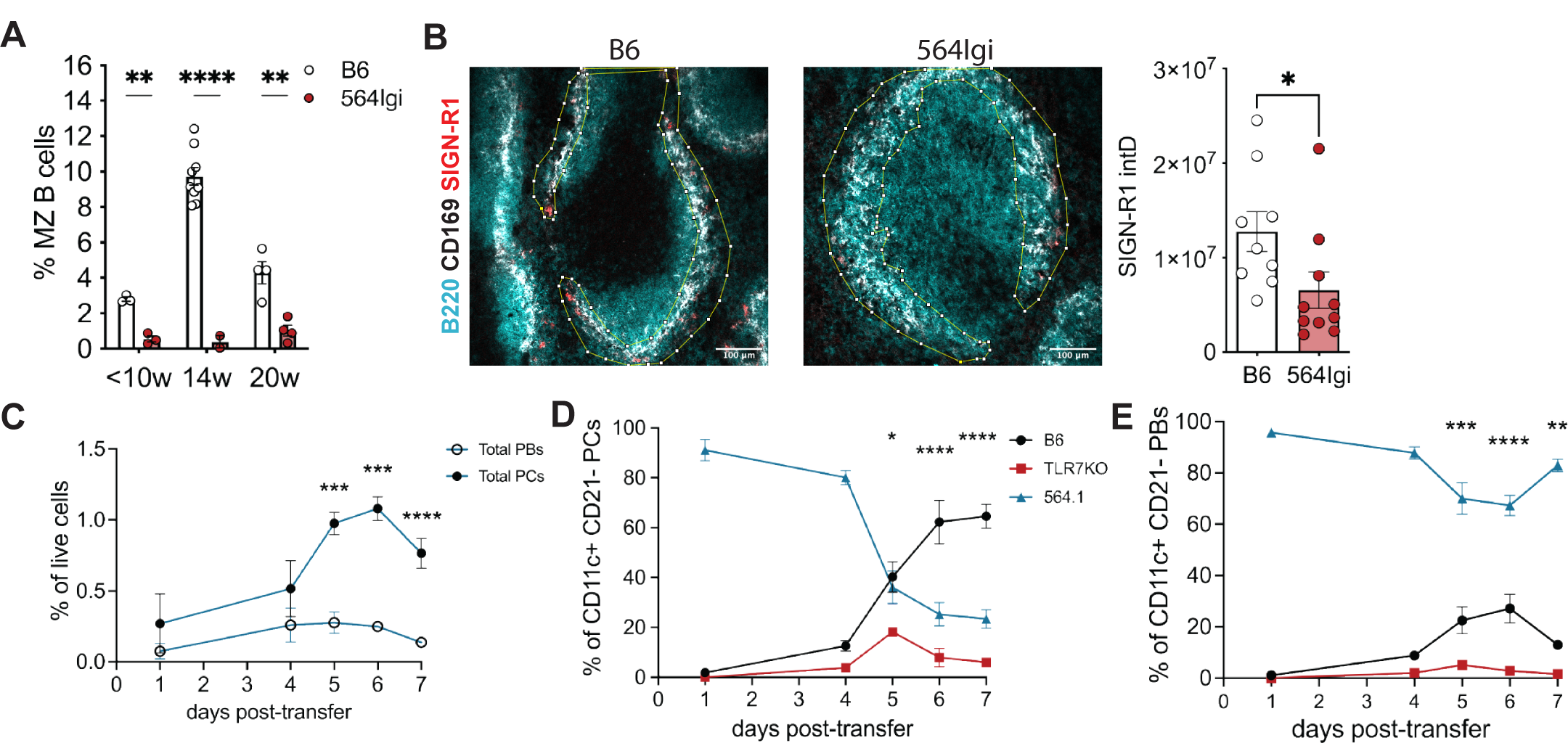
WT and *TLR7*-KO donor B cells occupy distinct splenic follicular niches and have divergent outcomes. related to Figure 2. (A) Frequencies of marginal zone B cells in naive B6 and 564Igi mice from flow cytometry analysis of splenocytes. <10w: n=3, 14w B6: n=9, 14w 564Igi: n=2, 20w B6: n=4, 20w 564Igi: n=4. (B) Representative IF images of naive B6 (n=9) and 564Igi (n=10) splenic follicles at 30x magnification with ROI marking the marginal zone and calculated SIGN-R1 integrated density (intD) within MZ ROI. Analysis completed from IF images of splenic follicles. Statistical analyses with Welch’s t test. (C) Frequencies of total plasma cells (PCs) and plasmablasts (PBs) over time course of 7 days from competitive transfer of WT and TLR7KO B cells. (D and E) Respective frequencies of WT B6, TLR7KO, and 564.1 CD11c+ CD21- PCs and PBs over competitive transfer time course of 7 days. Statistical analyses with Šidák’s multiple comparisons *=p<0.05,**=p<0.01, ***=p<0.001; ****=p<0.0001.

**Figure S3.**
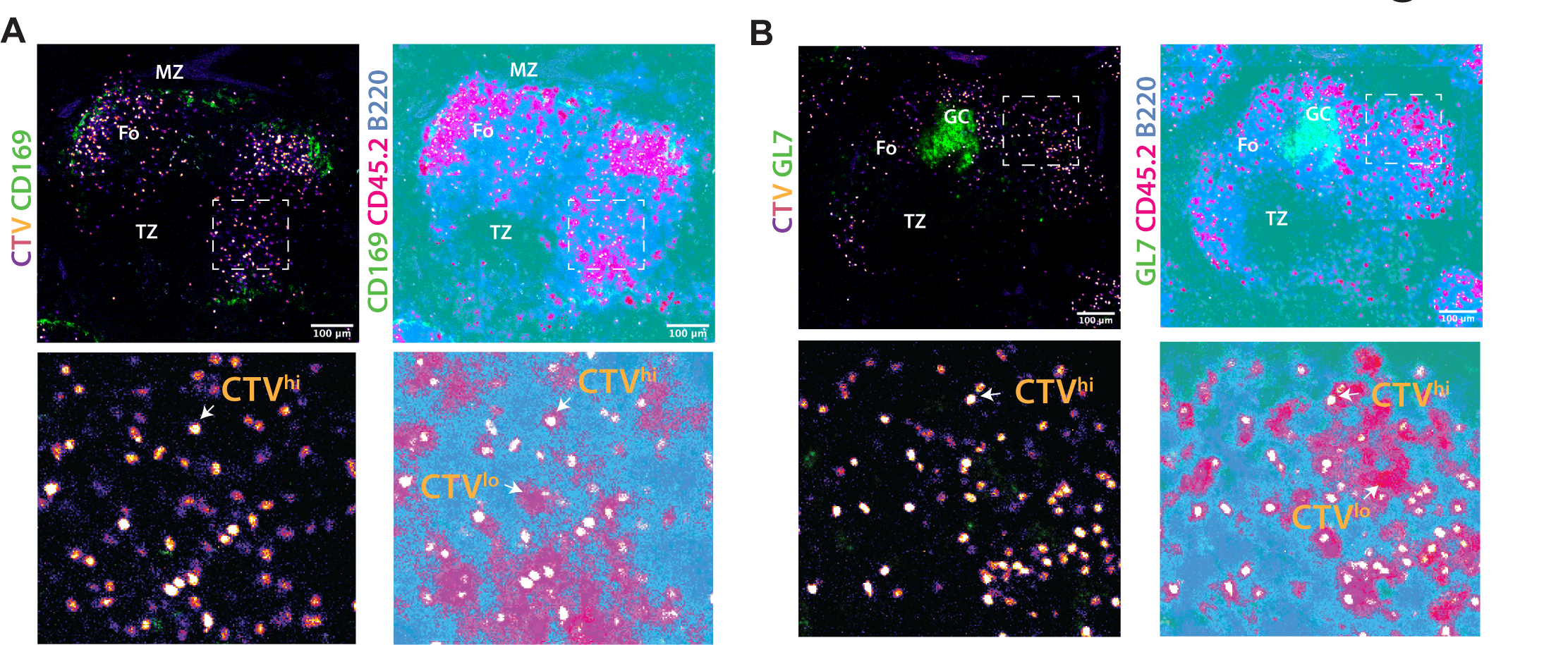
At 5 days post-transfer, donor B cells exhibit a fixed division program and require at least 7 divisions for ASC differentiation. Related to Figure 3. (A, B) IF imaging of 564Igi recipient spleens receiving CTV-labeled donor B cells with respect to marginal zone and germinal center. CTV-hi and CTV-lo donor cells are indicated with white arrows in enlarged regions of interest with white dotted borders. Image acquired at 30x magnification. Fo = follicle, TZ = T cell zone, MZ = marginal zone, GC = germinal center. (A) Representative flow cytometry plots showing CTV histograms of WT donor B cells and distrubution of CD21lo CD23-, CD21mid CD23+, and CD21+ CD23- subsets within resting (div. 0), dividing (div. 1-6), and terminally divided (div. 7+) donor cells. Quantification shows subset frequencies for both B6 and TLR7KO donors. (B, C) Respective frequencies and paired analysis of B6 and TLR7KO CD21+ CD23- and CD21mid CD23+ donor B cells with 0, 1-6, and 7+ divisions. Statistical analyses with paired t test, with p values indicated on plots. (D) Representative flow cytometry plots of CD11c versus CD138 for donor B6 and TLR7KO B cell subsets and frequencies of CD11c+ CD138- donors within specified subsets (n=5). Statistical analyses with multiple paired t test. *=p<0.05, **=p<0.01, ****=p<0.0001.

**Figure S4.**
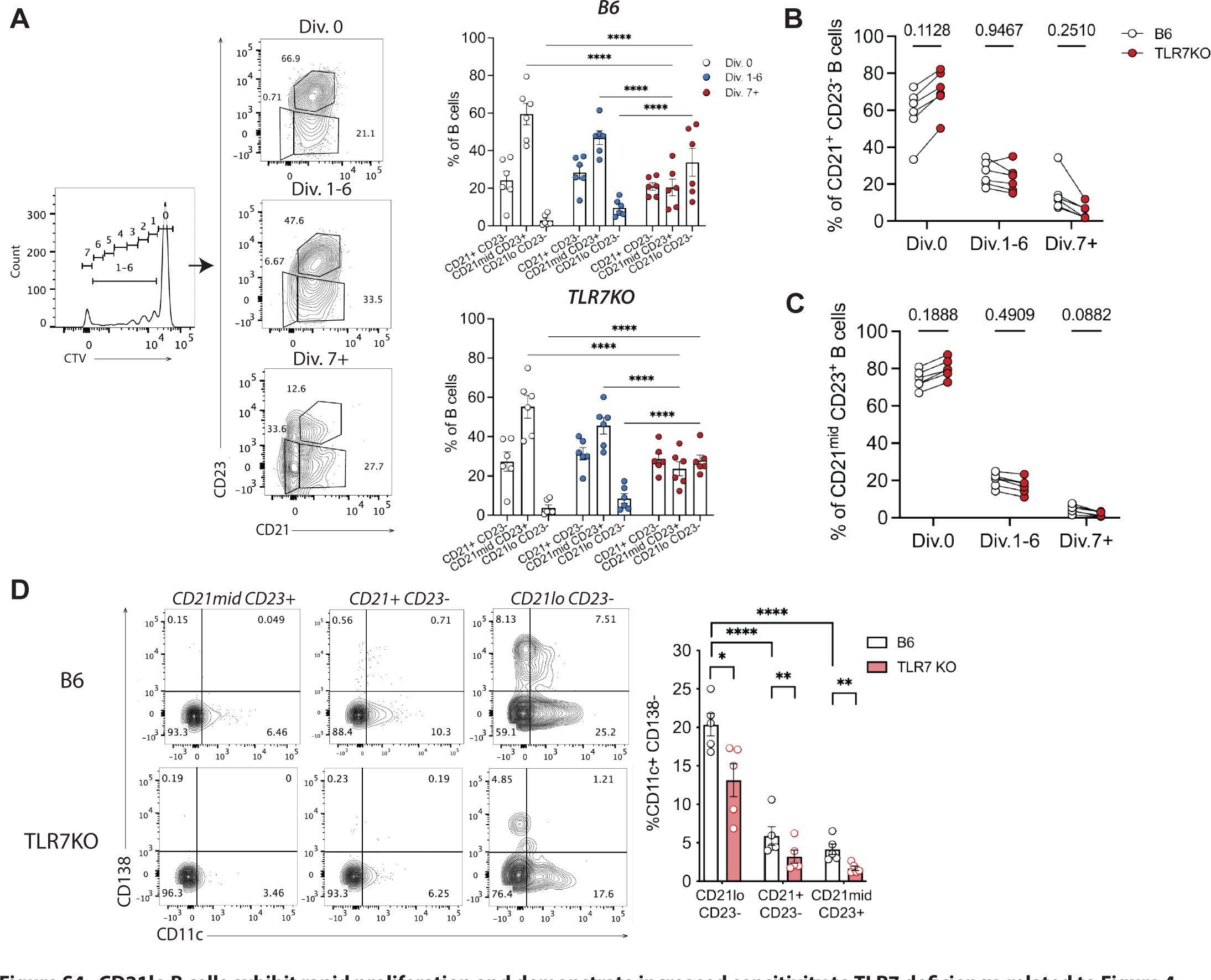
CD21lo B cells exhibit rapid proliferation and demonstrate increased sensitivity to TLR7 deficiency. related to Figure 4.

**Figure S5.**
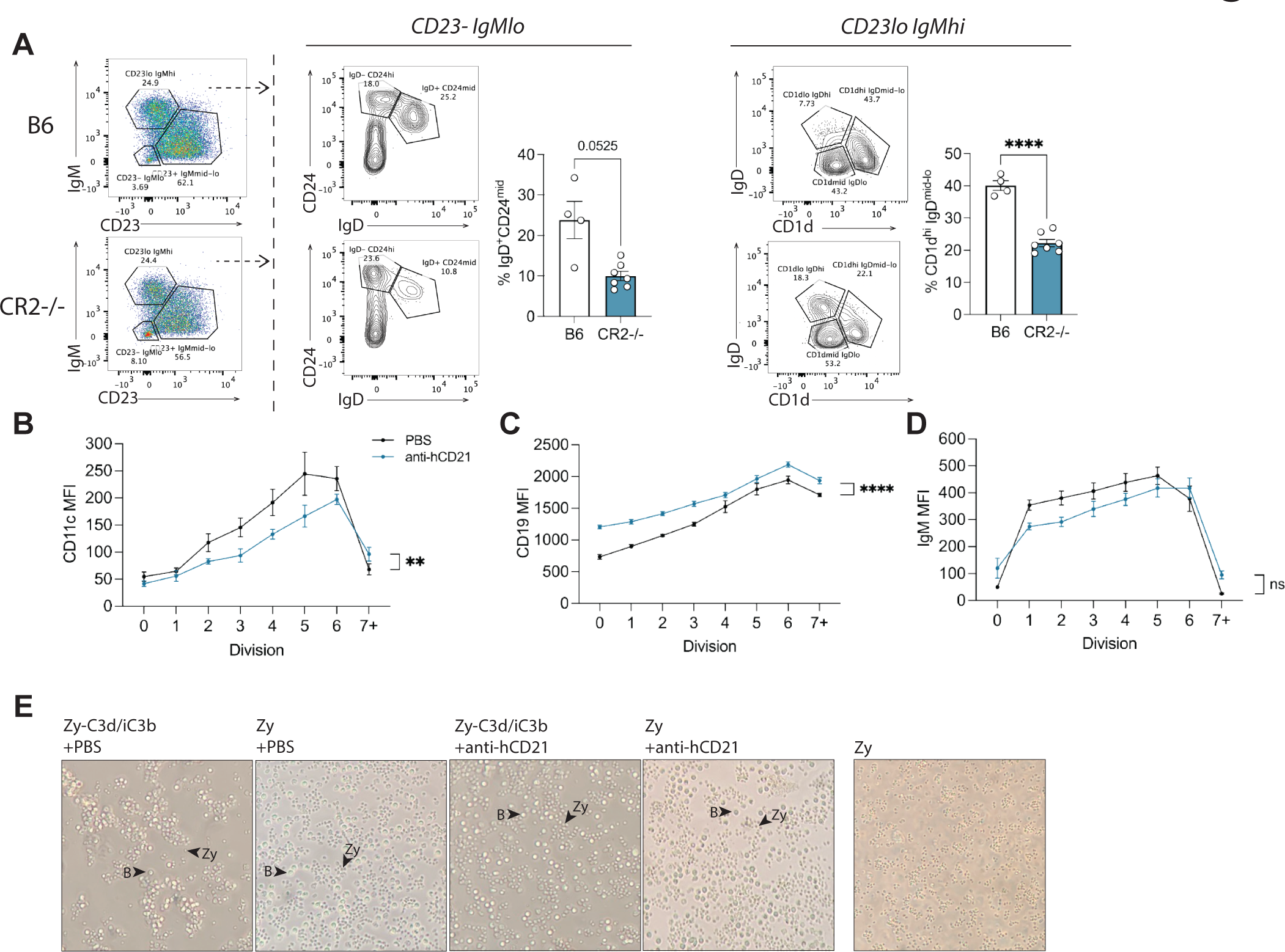
CD21 downregulation during proliferation is reversible and can be restored by functional blockade of the receptor. related to Figure 5. (A) Representative flow cytometry plots and corresponding quantifications of WT B6 (n=4) and CR2-/- (n=7) B cell compartments post- competitive transfer. (B) Surface CD11c MFI kinetics of CD21lo CD23- donor B cells per division treated with PBS or anti-hCD21 blocking antibody. (C, D) Surface CD19, and IgM MFI kinetics of total donor B cells per division treated with PBS or anti-hCD21 blocking antibody. (E) Representative bright-field microscopy images of B cells from *in vitro* hCD21 blockade at 20x magnification at 6 hr post-treatment. Image of Zy beads alone was taken as control. B cells and Zymosan beads are indicated with black arrowheads. B = B cells. Statistical analyses with area under the curve (b, c, d) and Welch’s t test (a-d). **=p<0.01, ***=p<0.001; ****=p<0.0001. ns = not significant.

**Figure S6.**
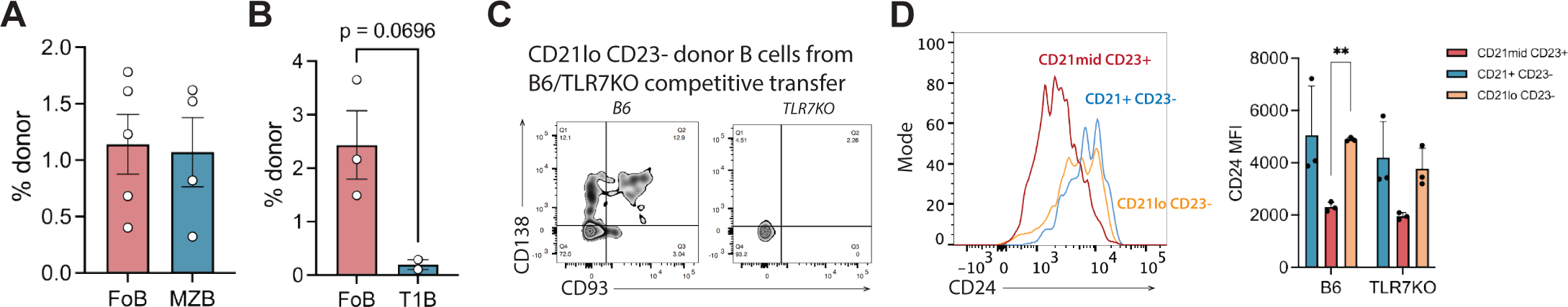
Follicular B cell derived CD21lo B cells are immediate precursors to autoreactive EF ASCs. related to Figure 6. (A) Frequencies of donor ASCs in host at 5.5 days post-transfer of sorted FoB and MZB. (B) Frequencies of donor ASCs in host at 5.5 days post-transfer of sorted FoB and T1B. (C) Representative flow cytometry plots of CD93 versus CD138 for B6 and TLR7KO CD21lo CD23- donor B cells (n=3). (D) Representative flow cytometry histograms of CD24 for CD21/CD23 subsets and corresponding quantification of CD24 MFI from B6 and TLR7KO donors at 7 days post-transfer (n=3). Statistical analysis with multiple paired t test, **=p<0.01.

**Figure S7.**
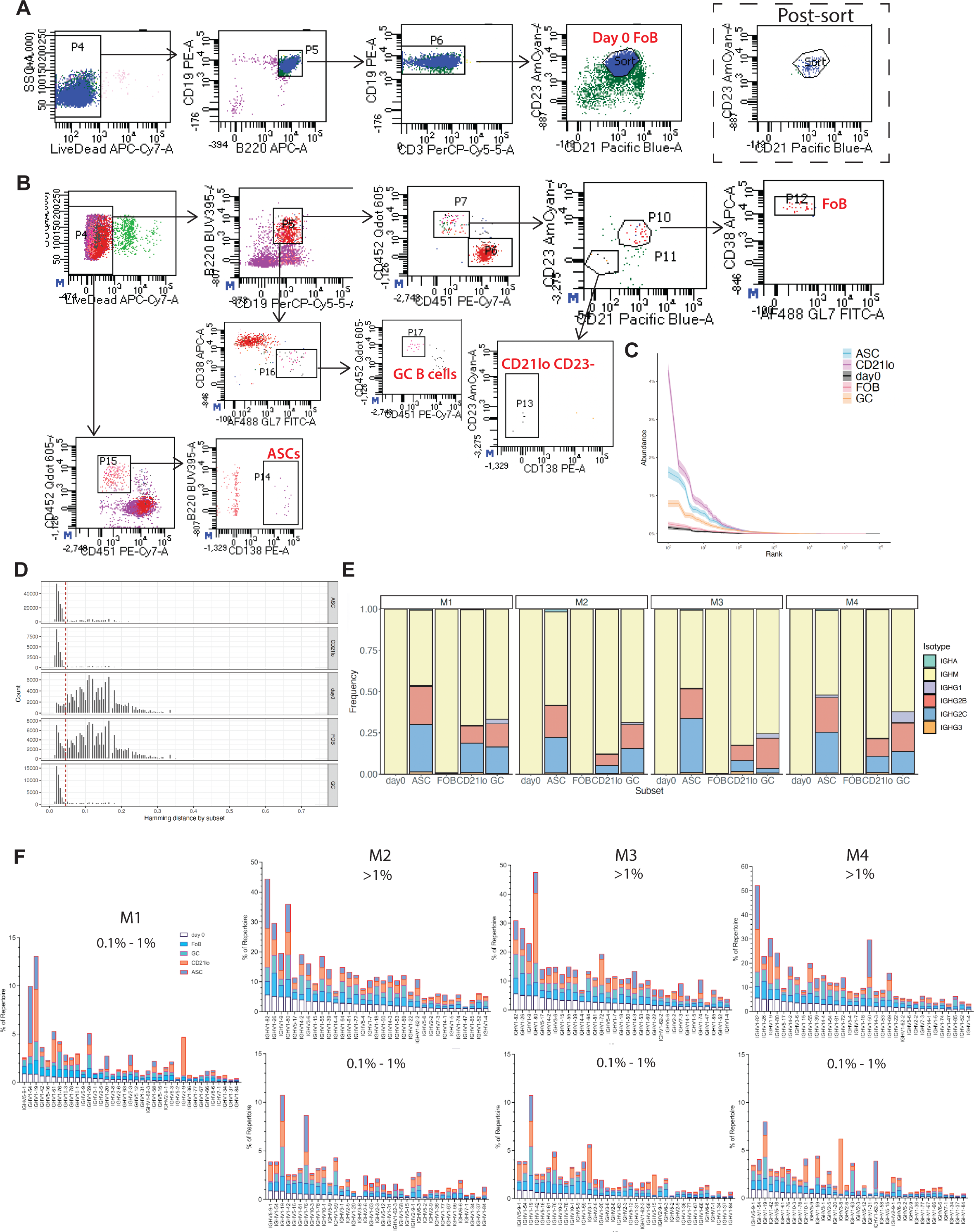
B cell repertoire sequencing reveals developmental trajectories of follicular B cells toward autoreactivity. related to Figure 7. (A, B) Respective sorting strategies of day 0 follicular B cell donors and progeny subsets at 6.5 days post-transfer. Sorted populations indicated in red. (C) Clonal abundance distribution curve of donor B cell subsets. Shadowed areas indicate 95% CI. ASC = ASCs, CD21lo = CD21lo cells, day0 = day 0 follicular B cells, FOB = day 6.5 follicular B cells, GC = GC B cells. (D) Hamming distance histograms by B cell subset. Red dotted line indicates threshold of 0.045 used for clonal clustering. (E) Plot of Ig isotype frequencies for each donor subset from individual mice (M1-M4). (F) Vh gene repertoire frequency plots of individual mice (M1-4). IGHV families making up to >1% and 0.1% - 1% of day 0 follicular B cell repertoire were plotted and respectively indicated.

**Table S1.**
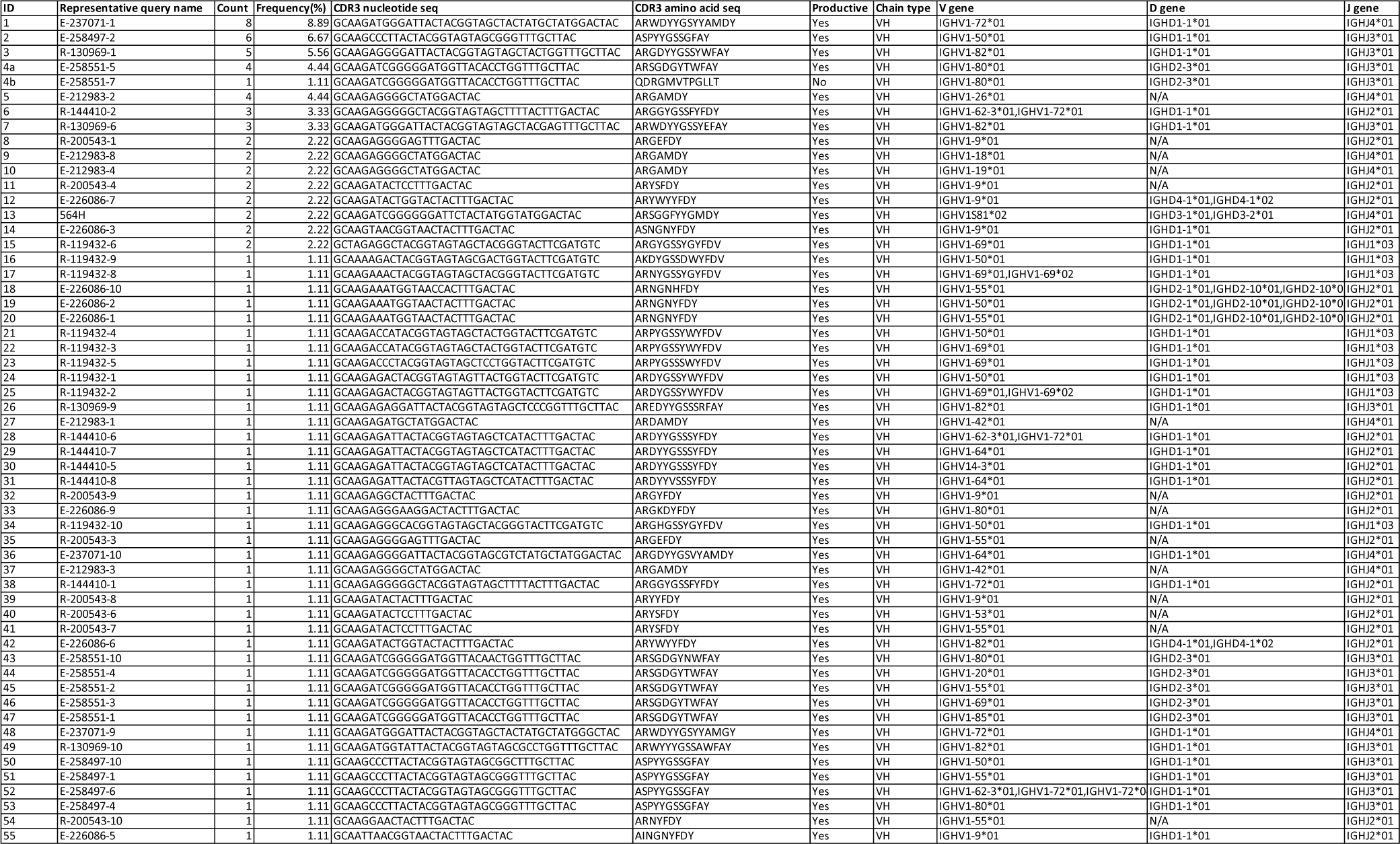
IgBLAST summary table of selected clones and sequences.

**Table S2.**
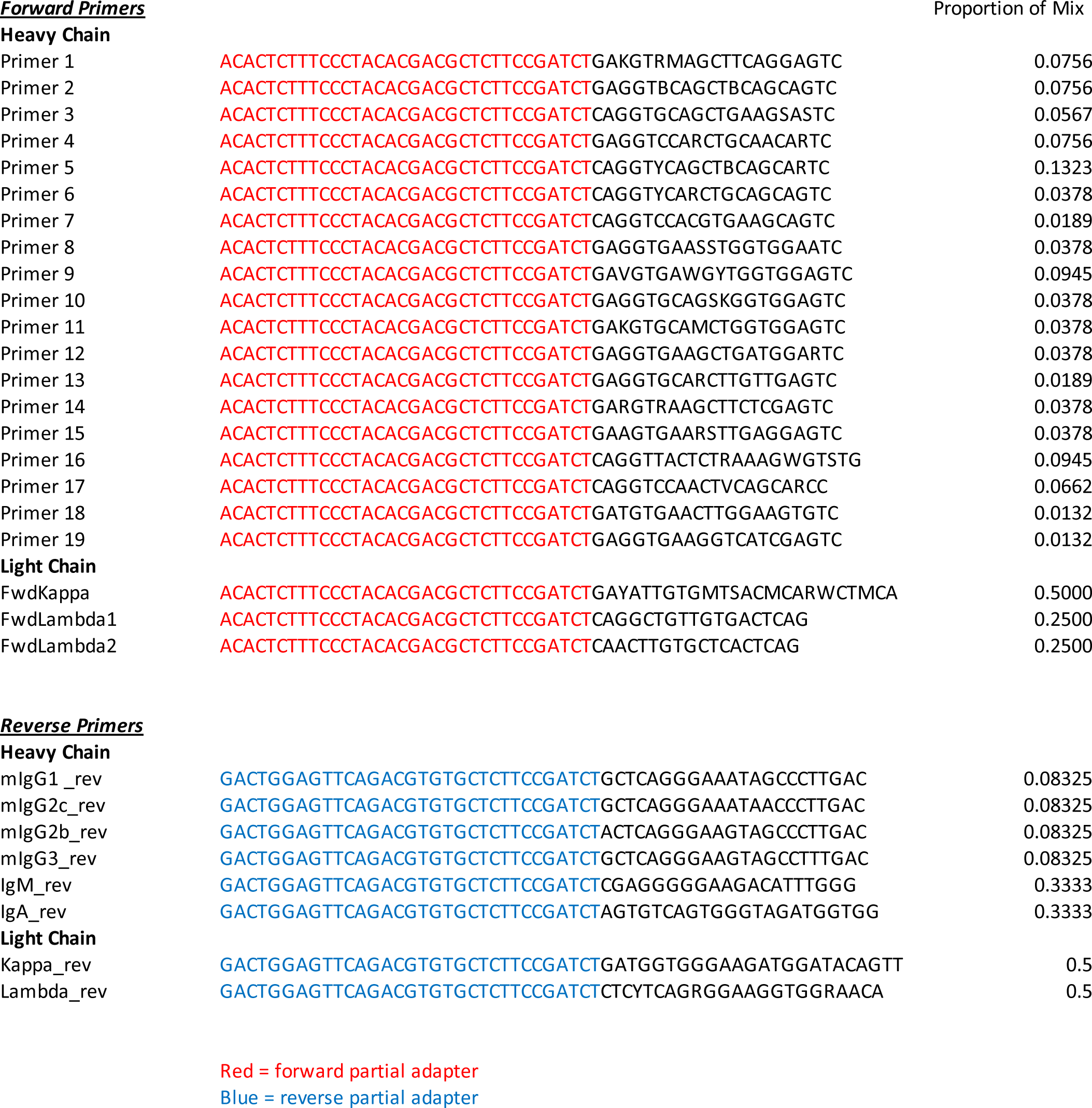
Primer sequences for BCR-seq library preparation.

